# Error-corrected deep targeted sequencing of circulating cell-free DNA from colorectal cancer patients for sensitive detection of circulating tumor DNA

**DOI:** 10.1101/2023.12.22.573042

**Authors:** Amanda Frydendahl, Mads Heilskov Rasmussen, Sarah Østrup Jensen, Tenna Vesterman Henriksen, Christina Demuth, Mathilde Diekema, Henrik J. Ditzel, Sara Witting Christensen Wen, Jakob Skou Pedersen, Lars Dyrskjøt, Claus Lindbjerg Andersen

**Affiliations:** Department of Molecular Medicine, Aarhus University Hospital, Denmark; Department of Clinical Medicine, Aarhus University, Denmark; Institute of Molecular Medicine, University of Southern Denmark, Denmark; Department of Oncology, Odense University Hospital, Denmark; Department of Oncology, Lillebaelt Hospital, Denmark; Bioinformatics Research Center, Faculty of Science, Aarhus University, Denmark

**Author notes:** **Corresponding author:** Claus Lindbjerg Andersen, Department of Molecular Medicine, Aarhus University Hospital, Palle Juul-Jensens Boulevard 99, Aarhus N, DK-8200, Denmark. These authors contributed equally (Amanda Frydendahl, Mads Heilskov Rasmussen).

## Abstract

**Introduction:** Circulating tumor DNA (ctDNA) is a promising biomarker, reflecting the presence of tumor cells. Sequencing-based detection of ctDNA at low tumor fractions is challenging due to the crude error rate of sequencing. To mitigate this challenge, we developed UMIseq, a fixed-panel deep-targeted sequencing approach, which is universally applicable to all colorectal cancer (CRC) patients.

**Methods:** UMIseq features UMI-mediated error correction, exclusion of mutations related to clonal hematopoiesis, a panel of normals for error modeling, and signal integration from single-nucleotide variations, insertions, deletions, and phased mutations. UMIseq was trained and independently validated on pre-operative (pre-OP) plasma from CRC patients (n=364) and healthy individuals (n=61).

**Results:** UMIseq displayed an area under the curve surpassing 0.95 for allele frequencies (AF) down to 0.05%. In the training cohort, the pre-OP detection rate reached 80% at 95% specificity, while in the validation cohort, it was 70%. UMIseq enabled the detection of AFs down to 0.004%. To assess the potential for detection of residual disease, 26 post-operative plasma samples from stage III CRC patients were analyzed. Detection of ctDNA was associated with recurrence (p =0.08).

**Conclusion:** UMIseq demonstrated robust performance with high sensitivity and specificity, enabling the detection of ctDNA at low allele frequencies.

## INTRODUCTION

Circulating tumor DNA (ctDNA) has emerged as a powerful biomarker reflecting the presence of tumor cells, tumor burden, and patient outcome across multiple cancers, including colorectal cancer (CRC)^1,2^. Mutation-based tumor-informed sequencing methods are among the most widely used ctDNA detection approaches^3–7^. These methods can be categorized into two types: 1) the bespoke approach, where a unique assay targeting patient-specific mutations is designed for each patient^5,8,9^, and 2) the fixed approach, where the same assay, usually a cancer-specific capture panel, is applied to all patients^7,10,11^. The strength of a bespoke approach is a high ratio of tumor DNA markers to the number of analyzed genomic base pairs. However, this comes at the expense of having to design a new assay for each patient, and the prolonged turnaround time associated with this. Although a fixed approach often has a lower ratio of tumor DNA markers to the number of analyzed genomic base pairs, compared to a bespoke approach, it has the advantage of convenience and shorter turnaround times.

For variant calling in circulating cell-free DNA (cfDNA), most studies have focused solely on single-nucleotide variants (SNVs)^11–13^, a combination of SNV, insertions, and deletion (INDELs)^7^, or, more recently, phased variants^14^. Due to the higher complexity of phased variants, and possibly also INDELs and multi-nucleotide variants (MNV), these variants often have a low error rate, making them attractive targets for the detection of low-frequency variants^14^. Furthermore, incorporating several independent mutational classes into a ctDNA caller may enhance the tumorigenic signal and increase the sensitivity of ctDNA detection approaches.

Here, we present a tumor-informed fixed-panel strategy, termed targeted **u**ltra-deep **m**utation-**i**ntegrated **seq**uencing strategy (UMIseq). UMIseq leverages unique molecular identifiers (UMIs) for comprehensive error correction and employs a panel of normal (PON) for mutation-specific error modeling. To exploit the information from all the somatic tumor-specific mutations identified within the regions covered by the fixed panel, UMIseq integrates the signal from all mutations into a single measure. Bayesian inference is used to assess if this signal is higher than expected based on the PON, i.e., if the sample is ctDNA positive. UMIseq leverages information from various classes of mutations: SNVs, INDELs, MNVs, and phased variants. We conduct a rigorous assessment of the performance of the UMIseq approach using a comprehensive training and validation cohort design, including an evaluation of minimal residual disease (MRD) detection in post-operative (post-OP) plasma samples from stage III CRC patients.

## MATERIALS AND METHODS

### Study population

For this study, 383 CRC patients and 107 healthy controls were recruited and divided into a training cohort, a validation cohort, and a panel of normal (PON) control cohort. Two patients (0.52%) were excluded because none of the mutations in the tumor overlapped with the UMIseq capture panel. The training cohort included 37 healthy individuals and 126 stage I-III patients (stage I (pT2pN0cMO): n = 32, stage II (pT3-4pN0cM0): n = 59, stage III (pT1-4pN1-2cM0): n = 28, stage IV: n = 7). The validation cohort consisted of 24 healthy individuals and 209 stage I-III patients (stage I (pT2pN0cMO): n = 52, stage II (pT3-4pN0cM0): n = 108, stage III (pT1-4pN1-2cM0): n = 49), 29 patients with polyp cancers (pT1pN0cM0) tumors, and 17 patients with adenomas. Post-OP plasma samples (n = 26) from stage III patients with at least 30 months of radiological follow-up, or radiological confirmed local or distant recurrence were used for assessing the performance of UMIseq to detect MRD after operation. The PON consisted of plasma samples from 46 healthy individuals.

From all patients, tumor tissue, peripheral blood mononuclear cells (PBMCs), and an 8mL pre-operative (pre-OP) plasma sample were collected. Healthy individuals were anonymously recruited through the blood bank at Aarhus University Hospital or through the Danish colorectal cancer screening program. From healthy individuals, 8 mL of plasma was collected.

### Sample collection and processing

Details on tumor and blood sample collection, DNA extraction, library preparation, capture of target regions, and sequencing are described in **Supplementary Information**. In brief, tumor DNA and normal DNA from PBMCs were used to establish mutational profiles for each patient, as well as identify mutations associated with clonal hematopoiesis. For 12 patients diagnosed with synchronous tumors, we used the mutational profiles from both tumors in the ctDNA analysis. cfDNA from 8 mL plasma and normal DNA from each patient was analyzed by targeted sequencing using a custom panel based on hybridization capture, which included the most frequently mutated genomic regions observed in patients with CRC^15^ (15,465 bp, **Suppl. Table 1**). All samples (tumor, normal, and cfDNA) were sequenced by paired-end (2×150 bp) sequencing using the Illumina NovaSeq platform. Details can be found in **Supplementary information**.

### Plasma variant calling and estimation of circulating allele frequency

Variants were called in cfDNA and PBMC DNA using the Shearwater algorithm^16^. Shearwater models the counts on the forward and backward strands with a beta-binomial model to compute a Bayes factor for each possible SNV and uses multiple samples to estimate local error rates and dispersion. Here, the implementation was abstracted to also include INDELs, complex MNVs, as well as phased mutations (**Suppl. Table 2**). To this end, every mutation, regardless of its class, was represented as a read count vector of length four, consisting of forward and backward strand alternative counts (ALT and alt), and forward and backward non-alternative counts (^ALT and ^alt). For mutations spanning more than one nucleotide (most INDELs and MNVs), the mean counts across the locus were used. The ALT/alt and ^ALT/^alt counts were extracted from the UMI consensus BAM files. For both PBMC and cfDNA samples, the analyses were tumor-informed, i.e., counts were only extracted for mutations reported in the tumor VCF file of the matching patient. The bbb function from the R package deepSNV^17^ was used to calculate a mutation score, *m,* for each mutation, which equals the Bayes factor generated by bbb using the parameters model = ‘AND’ and prior = 0.5. The plasma samples of the PON cohort were processed in parallel with all other plasma samples and used to estimate the dispersion rho. The product of the Bayes factors of each sample’s mutation catalog was calculated to generate an integrated score, *s*, representing the likelihood of the sample being negative over the likelihood of it being positive. To mitigate the possibility that the noise structure of a particular sample deviated significantly from that of the PON, the integrated score was transformed into the UMIseq score, S. The UMIseq score represents a one-sided empirical p-value expressing how extreme the observed score *s* is compared to all other possible *s*-scores that could be generated in the sample using n mutations. To do this, a distribution of sample scores **S^n^**was generated from K random scores, i.e., sample scores from random mutation catalogs of length n sampled from all possible positions of the panel. The UMIseq score S was then calculated as the rank of the observed score *s* in **S^n^**, specifically S = 1 - (r+1)/(K+1), where K = 100,000, and r is the number of random scores larger than *s*. Plasma samples with a score above a fixed threshold resulted in the sample being called ctDNA positive. As an estimate of the mutational burden observed in a plasma sample, the mutational cAF was calculated as the sum of reads supporting the set of patient-specific tumor-informed mutations, divided by the sum of all reads across the mutated loci.

### Blacklisting of SNVs and INDELs

A total of 69 mutations were blacklisted as they were recurrent in plasma cfDNA from healthy individuals. Details can be found in **Supplementary Information**.

### Flagging of mutations associated with clonal hematopoiesis

UMIseq analysis of PBMC samples matched to each patient was performed to explore if any of the mutations identified in the tumor, also co-occurred in hematopoietic cells (Clonal hematopoiesis of indeterminate potential, CHIP). Details on this can be found in **Supplementary Information**.

### Limit of detection calculation

The limit of detection (LOD) of individual SNVs and INDELs within the capture panel was assessed to explore differences in LOD. A distribution of sample UMIseq LODs was similarly estimated. Details can be found in **Supplementary Information**.

### In silico estimation of the ctDNA detection probabilities

Details on *in silico* estimation of ctDNA detection probabilities at various cAFs can be found in **Supplementary Information.**

### Recurrent COSMIC mutations

Details on identification of recurrent COSMIC mutations within the UMIseq capture panel can be found in **Supplementary Information**.

### Analytical sensitivity analysis using a synthetic mixture

Details on assessment of the analytical sensitivity can be found in **Supplementary Information**.

### Tumor-informed model training of the UMIseq algorithm

To generate a UMIseq score threshold for calling a plasma sample ctDNA positive, UMISeq was applied to a training cohort consisting of pre-OP plasma from CRC patients (n = 126) and healthy controls (n = 37). To utilize the full diversity of the healthy control samples and the training cohort mutation catalog (n = 276 unique mutations), 25 Monte Carlo simulations were made as follows. UMIseq scores S from 100 randomly selected CRC cfDNA samples were calculated using their matching tumor catalog and used as positive labels. One hundred UMIseq scores representing non-cancer samples (negative labels) were calculated by applying *in silico* generated mutation catalogs to randomly selected healthy control cfDNA samples. Each *in silico* catalog was generated by sampling n mutations without replacement from the 276 mutations in the training cohort with probabilities equal to their relative frequency. The number of mutations (n) to sample in each catalog followed the distribution of mutation per patient in the training cohort. For each of the 25 simulations, the receiver operator characteristic (ROC) statistics, as well as the UMIseq S score resulting in a 5% FPR (corresponding to 95% specificity), were calculated. The mean UMIseq score of the 25 simulations was then used as a fixed cutoff, x, for calling plasma samples ctDNA positive in the validation cohort, and in the orthologous method, reproducibility, and repeatability tests, as well as for estimating the LOD of UMIseq.

### Assessment of UMIseq robustness

For a subset of samples, the robustness of UMIseq was assessed by i) comparison to an orthogonal method (ddPCR), ii) procuring, processing, and analyzing a new plasma aliquot, and iii) by generating two cfDNA libraries from the same cfDNA sample. Details can be found in **Supplementary Information**.

### Validation of specificity

Circulating cell free DNA from 24 plasma samples from healthy controls was extracted and subjected to UMIseq. These were used as independent controls to validate the 95% specificity that was set in the model training. We repeated the procedure for generating negative sample sets as described for the training cohort (see above). Random mutation sets (n = 100) were sampled from the training cohort tumor catalog according to their individual frequencies and used to generate 100 UMIseq scores S in the 24 validation controls. The specificity was calculated as the fraction of scores resulting in a negative call. This was repeated 25 times, and a t-test applied to test the difference in mean specificity between validation and training.

### Statistical considerations and calculations

Statistical tests were used as appropriate and as described in the text. All calculations were done with R (3.5.1) in the Rstudio environment (1.1.456) and with packages deepSNV^17^ (1.28.0), genomicranges (1.34.0), rsamtools (1.34.0), abind (1.4.5), dplyr (1.4.3), survival (3.1) and reshape2 (1.4.4) installed under the conda (23.3.1) environment.

## RESULTS

### The UMIseq method and study design

We first evaluated UMIseq (method overview in **Figure 1A**) by characterizing the general error profile across all positions in the capture panel, and then by assessing the analytical performance using a set of synthetic mixture samples with varying concentration of tumor DNA mixed in normal DNA (**Figure 1B**). Next, we trained UMIseq on a cohort of plasma from healthy individuals (n = 37) and pre-OP plasma samples from patients with stage I-IV CRC (n = 126). The performance of UMIseq was validated in an independent cohort of pre-OP plasma samples from patients with stage I-III CRC (n = 209), minimally invasive pT1pN0 cancers (n = 29), and non-invasive colorectal adenomas (n = 17) as well as plasma from healthy individuals (n = 24) (**Figure 1B**). Finally, the utility of UMIseq to predict minimal residual disease was assessed in post-OP samples from stage III CRC patients (n = 26) (**Figure 1B**). UMIseq of plasma samples resulted in median UMI consensus sequencing depths of 8367 (IQR 9225) and 10353 (IQR 9191) for the training and the combined validation and MRD cohorts, respectively (**Suppl. Figure 1A**). Of note, the median sequencing depths are partly determined by the cfDNA input. The median plasma cfDNA conversion efficiency was 88% (IQR 47%) and 83% (IQR 52%) for the training and the combined validation and MRD cohorts, respectively (**Suppl. Figure 1B)**. Comparable median depths were obtained from UMIseq on PBMC samples used for CHIP filtering (**Suppl. Figure 1C-D**). The clinicopathological characteristics of the training and validation can be seen in **Table 1**. The applied capture panel was designed to capture at least one mutation in all patients. In practice, 93% (353/381) of patients had at least two mutations within the capture panel (median 3 mutations per patient, IQR 2) (**Table 1, Suppl. Figure 2**).

**Figure 1.**
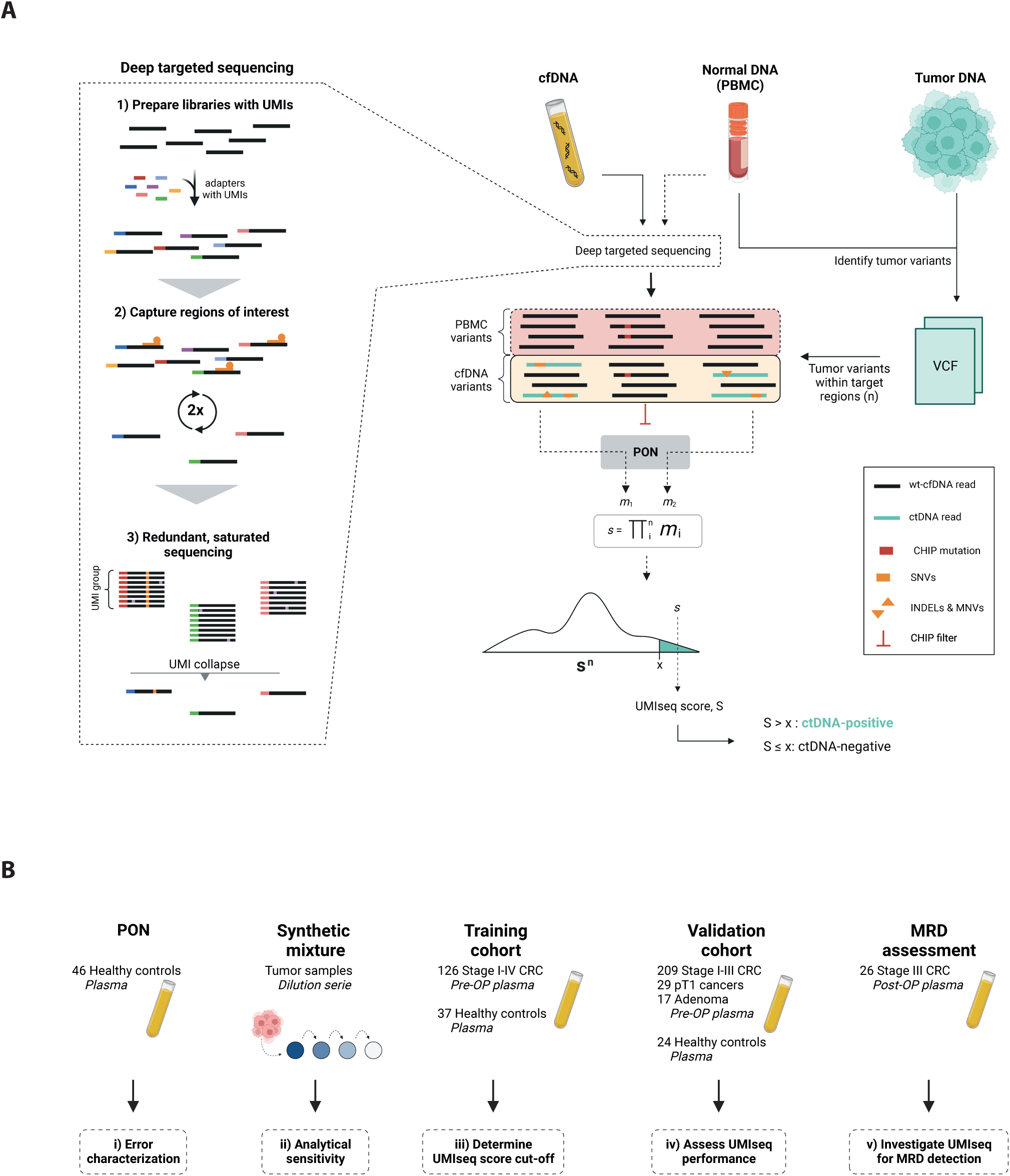
Study overview. **A)** Left panel: The workflow for deep target sequencing. (step 1) library preparation using UMI-containing sequencing adapters, (step 2) two consecutive rounds of hybridization-based capture of genomic regions frequently mutated in CRC, (step 3) redundant, saturated sequencing to ensure the creation of UMI families with at least 3 reads, each of which are finally collapsed into a consensus read, thereby eliminating random sequencing errors. Right panel: cfDNA from plasma, and optionally also DNA from PBMCs, are subjected to deep targeted sequencing. DNA extracted from the tumor and PBMCs are used to identify tumor-specific variants (n), which are then searched for in the consensus reads from deep-targeted sequencing of cfDNA and PBMC DNA. Variants present in the PBMC DNA are classified as CHIP mutations and excluded. For each of the n variants observed in cfDNA a mutation score, *m*, is calculated based on the noise and dispersion observed in the panel-of-normal (PON). By integrating the signal from all *m* scores, a sample score, *s*, is calculated. To mitigate the possibility that the noise structure of a particular sample may deviate significantly from that of the PON, the integrated sample score *s* is transformed into the UMIseq score S. To do this, a sample-specific score distribution, **S^n^**, is generated by calculating UMIseq scores in the cfDNA sample from 100,000 in silico random mutation catalogs harboring the same number of mutations as in the sample. S is then calculated as the rank of the integrated score of the tumor mutation catalog in the distribution **S^n^**. Samples with a UMIseq score above a fixed threshold, x, are deemed ctDNA-positive. **B)** Overview of the cohorts used in this study. Figure created with Biorender.com.

**Table 1.**
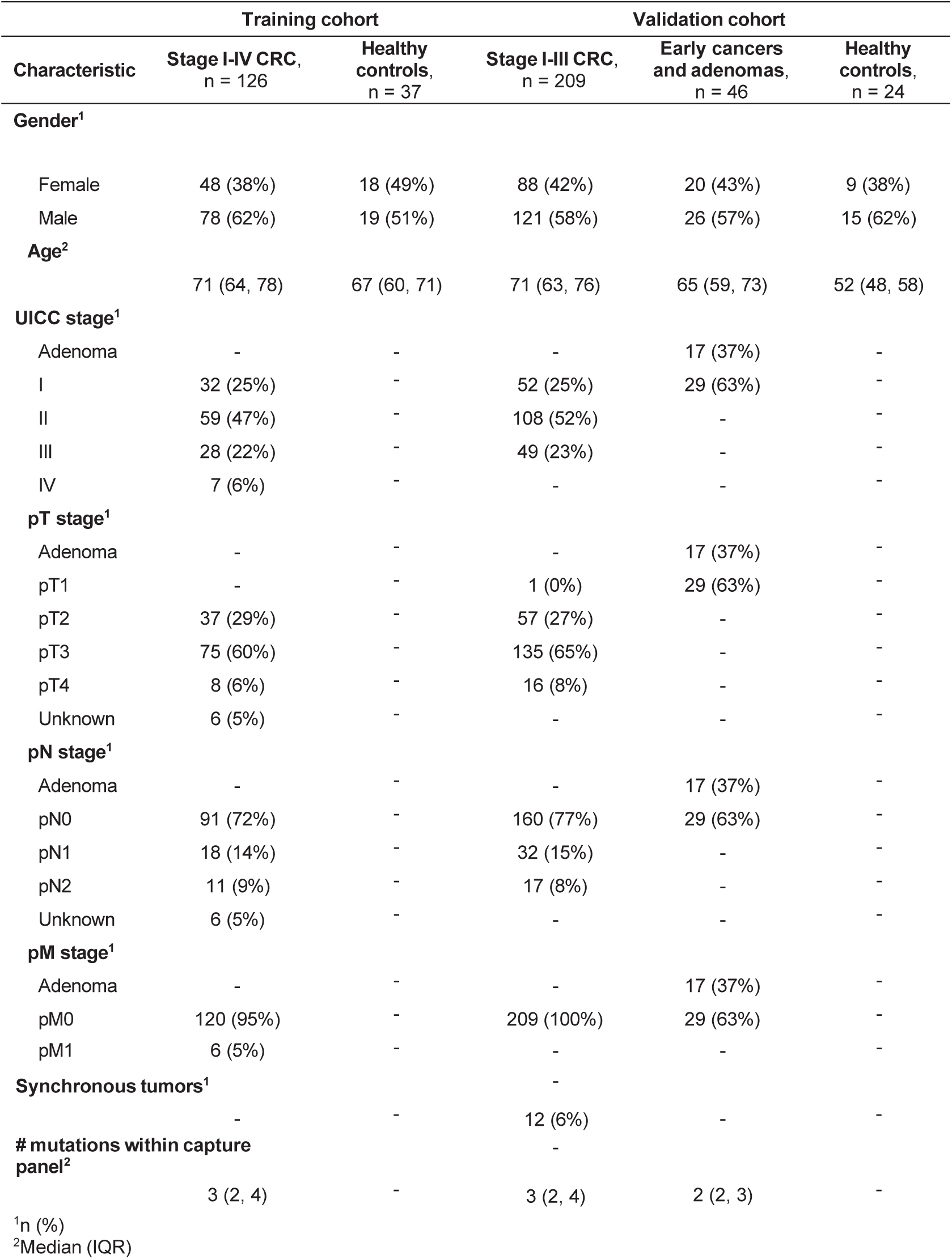
Cohort characteristics.

### Error characterization of UMIseq

To form a PON, UMIseq was applied to cfDNA from healthy individuals (n = 46) and sequenced to a median depth of 9086 (IQR 5464) (**Suppl. Figure 1A**). As the PON samples are not expected to carry any cancer related mutations, it facilitated an exploration of the error profile associated with UMIseq. All non-reference base calls at each genomic position targeted by the capture panel were considered errors. The absolute number of errors in each plasma sample was positively correlated with sequencing depth (r = 0.79; p < 0.0001, Pearson’s correlation). The most frequently occurring errors were G>T transversions, and C>T and G>A transitions (**Figure 2A**). More than 90% of the C and G sites within the panel had C>T, G>T, and G>A substitutions with an error rate above 0.001%, and for approximately 25% of the sites, the noise was higher than 0.01% (**Figure 2B**). In contrast, between 75% and 80% of T>G, A>C, C>G and G>C substitutions had an error rate of less than 0.001% (**Figure 2B**). Deletions, and in particular insertions, displayed low error rates (**Figure 2B**). Further evaluation including the trinucleotide context of each substitution revealed that the error rate of certain substitutions was highly affected by the neighboring 3’ or 5’ nucleotides, while others were not. Regardless of the trinucleotide context, T>G and C>G substitutions generally had a low error rate, whereas C>T substitutions had the highest error rates. The error rates of C>T substitutions were particularly high when the 3’ base was G, i.e. N(C>T)G, independent of the 5’ base (**Figure 2C**).

**Figure 2.**
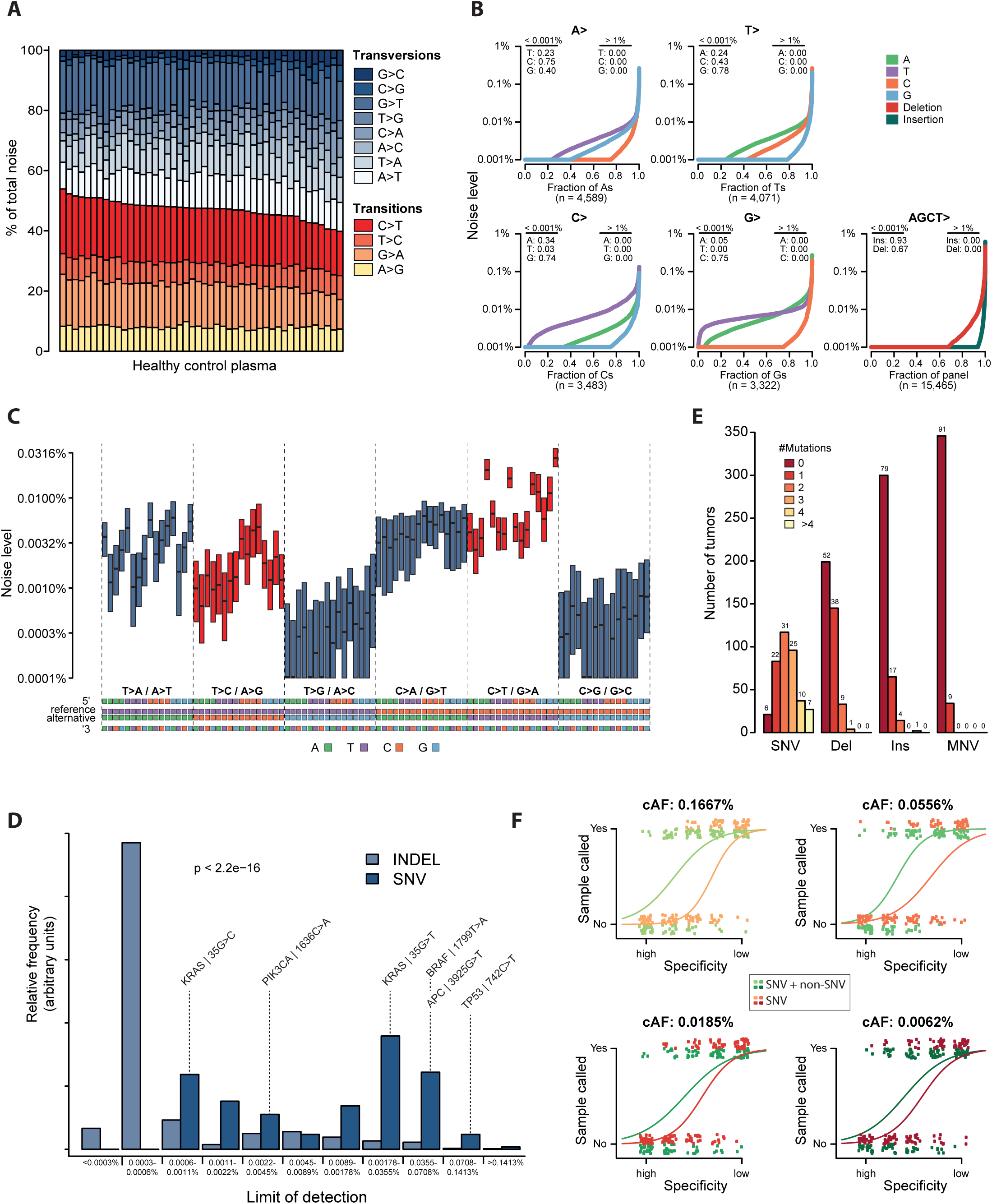
Error characterization of target regions. **A)** Noise distribution shown as relative frequency of single nucleotide errors observed in healthy control plasma (n = 46) across the target regions (15,465 bp). **B)** The mean noise across healthy plasma controls for each single-base substitution, insertion, and deletion. The fraction of sites with a noise level below 0.001% and above 1% are indicated. **C)** Boxplot of the noise level (median and IQR) of single-base substitutions according to their trinucleotide context. **D)** Predicted limit of detection (LOD) of SNVs (dark blue) and INDELS (light blue) at 99% specificity. The LODs of common CRC mutations are annotated. **E)** The frequency of SNVs, deletions, insertions, and multi-nucleotide variants (MNV) in tumors from all patients (n = 381). The fraction (in percentage) is indicated at the top of each bar. **F)** Simulated probabilities of calling a sample positive with or without non-SNVs alterations at various circulating allele frequencies and specificities.

### Assessment of the theoretical limit of detection

Based on the error rates observed when applying UMIseq to cfDNA from healthy individuals, we predicted the LOD for SNVs and single-base INDELs. This revealed that INDELs had a significantly different, and generally lower, LOD distribution compared to SNVs (p < 0.0001, Kolmogorov-Smirnoff test) (**Figure 2D**). Notably, 48% of CRC patients exhibited at least one deletion and 21% had at least one insertion within the capture panel (**Figure 2E**). In addition, MNVs were observed in 9% of CRC patients. To assess the sensitivity of UMIseq based solely on SNVs or a combination of SNVs, INDELs, and MNVs, we generated 25 *in silico* patient mutational catalogs by random selection of SNVs and non-SNVs (INDELs and MNVs) from the collective mutational catalog of all patients in the study (as described in **Suppl. Information**). Next, we generated synthetic samples for each *in silico* patient, by Poisson sampling 20,000 counts using a Poisson rate λ = cAF for each mutation. This process was carried out at four distinct cAFs: 0.1667%, 0.0556%, 0.0185%, and 0.0062%. For each patient, two integrated *s*-scores were generated: one including both SNV and non-SNV mutations and one including only the SNV mutations (i.e., INDELs and MNVs were removed from the mutation catalog). Inclusion of the non-SNVs increased the likelihood of correctly assigning a ctDNA positive label to samples across all cAFs, with the tendency increasing with higher cAF (**Figure 2F**).

### Analytical sensitivity of UMIseq

To evaluate the analytical sensitivity of UMIseq, we mixed tumor DNA from 36 patients to create a compound sample (Sample A) with 95 ground truth mutations within the target regions of the capture panel (the foreground mutations) (**Figure 3A**). Sample A was serially diluted with normal PBMC DNA (Sample X) to create five synthetic ctDNA samples (Samples B-F) with decreasing tumor DNA fractions. Comparison of the error rate of the foreground mutations (n = 95) to that of recurrent CRC mutations (n = 275) occurring in the COSMIC database^17^ showed a great similarity (p = 0.2928, Kolmogorov–Smirnov test), indicating that the foreground mutations were representative of CRC mutations (**Figure 3B**). Of note, the error rate across the entire panel (background) was lower than both the foreground and the COSMIC mutations (p < 0.0001, Kolmogorov-Smirnoff test), which may hint at an increased underlying error rate of recurrent mutations in CRC.

**Figure 3.**
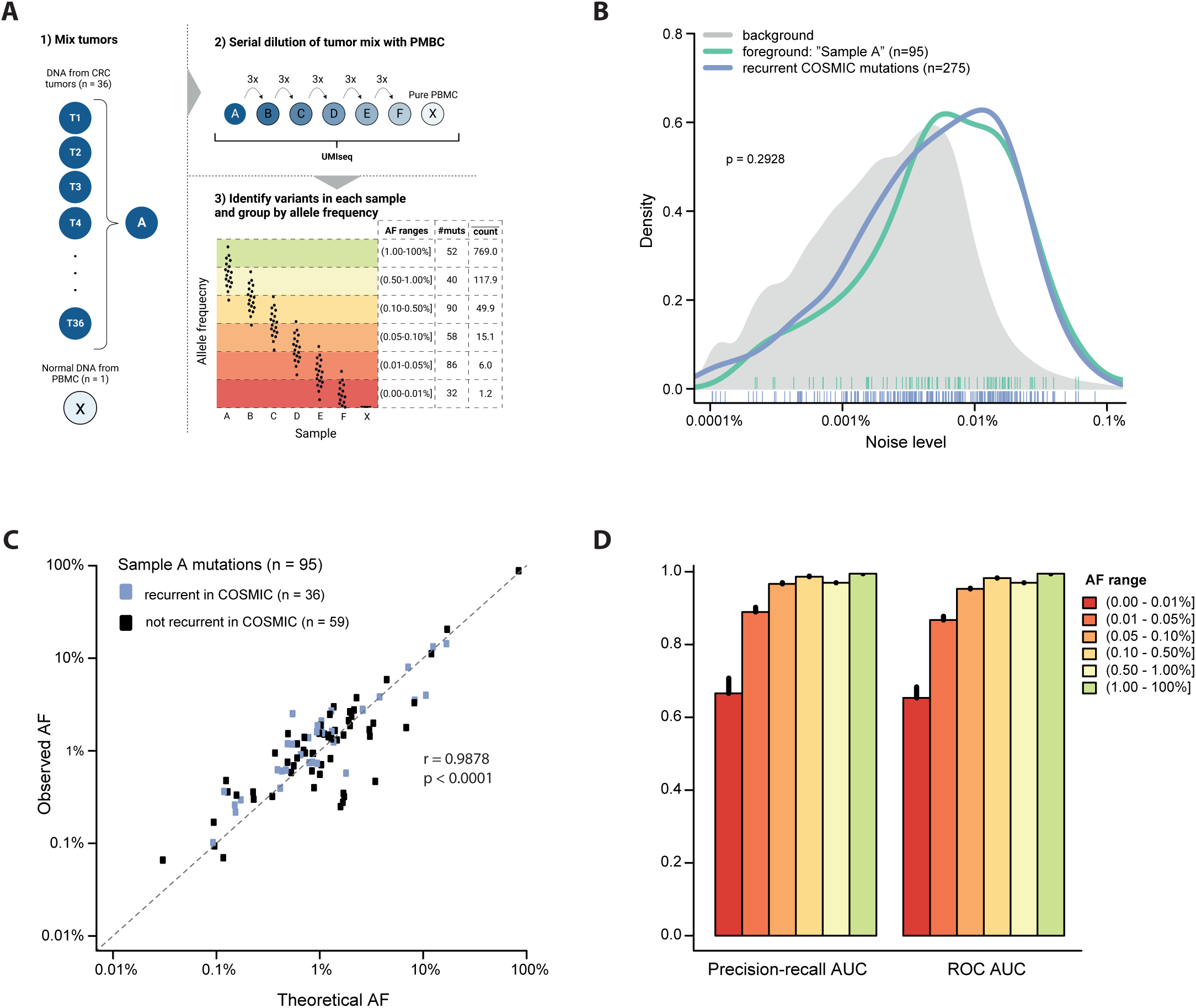
Analytical sensitivity performance. **A)** Schematic overview of the strategy for creating and analyzing artificial sample sets with known variants (foreground, n = 95 mutations identified in 36 tumor samples T1 through T36). The AF ranges, the number of mutations (#muts) and the mean number of reads supporting the mutations in the AF range (count) are indicated. Created with Biorender.com. **B)** The mean noise of foreground and recurrent COSMIC mutations in healthy plasma controls (n = 46) displayed as kernel densities with individual mutations indicated below. The density distribution of random substitutions is shown in grey. **C)** Correlation between theoretical AF (calculated from the individual tumors mutation AFs) and the observed AF in sample A of the foreground mutations. Mutations unique to sample A are shown in black (n = 59) while mutations reported to be recurrent in COSMIC are shown in blue (n = 36). **D)** AUC for precision-recall and ROC curves at six different AF ranges with error bars indicating the 95%CI on the estimation from 20 Monte-Carlo simulations.

There was a good correlation between the expected AFs of the foreground mutations in Sample A and the observed AFs from UMIseq (r = 0.9878, p < 0.0001, Pearson’s correlation) across an AF span from 0.1% to almost 100% (**Figure 3C**). The strong correlation between expected and observed AFs was independent of whether the mutations occurred in COSMIC (n = 36; r = 0.9934, p < 0.0001) or not (n = 59; r = 0.9011, p < 0.0001). By grouping the mutations observed in samples A through F based on their observed AF (**Figure 3A**), we assessed the performance of UMIseq as a function of AF. Remarkably, UMIseq displayed an area under the curve (AUC) above 0.95 for the precision-recall and ROC estimates down to AFs of 0.05% (**Figure 3D**). At AF below 0.05%, the absolute number of reads supporting the mutations approached a single read, *e.g.* the mean number of reads supporting each mutation was 1.2 reads in the AF range of 0-0.01%. This indicates that at very low AFs, sampling effects, rather than the error rate of UMIseq, determines whether ctDNA is detected.

### Training of UMIseq for ctDNA detection using pre-OP plasma

The UMIseq algorithm was trained to call ctDNA in plasma samples with 95% specificity using a cohort of pre-OP plasma samples from patients with stage I-IV CRC (n = 126; stage I: 32; stage II: 59; stage III: 28; stage IV: 7), and plasma from a control group of healthy individuals (n = 37). ROC curve analysis showed an AUC of 0.90, demonstrating that UMIseq has both high sensitivity and specificity (**Figure 4A**), and that integrating signal from all available mutations in UMIseq increases performance compared to a strategy where only a single mutation is used (ROC AUC 0.83, **Suppl. Figure 3**). We estimated the LOD of UMIseq as the minimum cAF required to call a sample ctDNA positive with 95% specificity (UMIseq score cutoff = 0.9797, used henceforth). An inverse correlation between LOD and sequencing depth was seen, with the median LOD reaching 0.011% (IQR 0.029%) and 10% of estimated LODs below 0.001% at 20,000x depth (**Figure 4B**). UMIseq detected ctDNA in 80% (101/126) of the pre-OP samples. The detection rate was 69% (22/32) in stage I, even higher in stage II (85%; 50/59) and stage III (79%; 22/28), and the highest in stage IV (100%; 7/7) (**Figure 4C**). The ctDNA detection rate increased with the pathological T (pT) category (pT2: 68%, pT3: 83%, pT4: 100%), while there was no correlation to the pathological N (pN) category (pN0: 79%, pN1-2: 79%) (**Figure 4D**).

**Figure 4.**
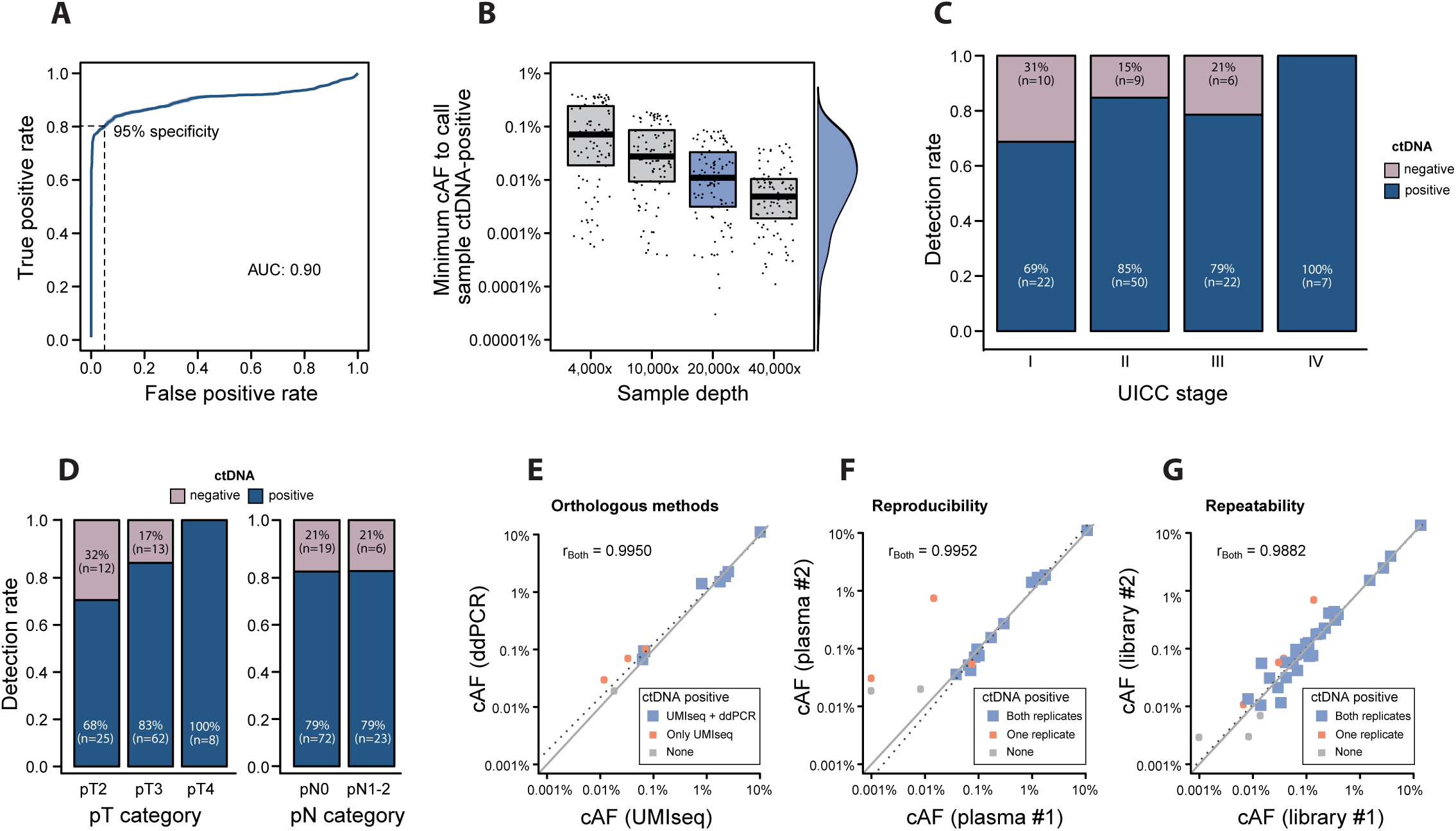
Training cohort. **A)** ROC analysis on pre-OP plasma samples for UMIseq. Pre-OP plasma samples (n = 126) were used as positive labels, and plasma samples from healthy controls against *in silico*generated mutational catalogs based on the patient tumors were used as negative labels. The UMIseq sample score threshold for ctDNA detection was set at 95% specificity, as indicated. **B)** Simulated sample LODs for UMIseq at various sequencing depths (4,000x, 10,000x, 20,000x, and 40,000x). The minimum cAF required to call a sample ctDNA-positive was calculated for n = 100 *in silico*-generated mutational tumor catalogues at four sequencing depths. The density (right) indicates the sample LOD distribution at 20,000x mean depth (blue). **C)** ctDNA detection rates stratified by UICC stage. **D)** ctDNA detection rates stratified by pT (left) and pN category (right). **E-G)** UMIseq robustness was evaluated by comparing ctDNA status and cAF levels in paired plasma samples purified from the same blood sample and analyzed with UMIseq and ddPCR (n = 11 blood samples) **(E)** or with UMIseq twice (n = 19 blood samples) **(F)**, and in paired cfDNA samples analyzed with UMIseq (n = 46 plasma samples) **(G).** Sample pairs where both, one or none of the paired samples were called ctDNA positive are shown in blue, red and grey, respectively. Using pairs with both samples ctDNA-positive, the Pearson’s correlation coefficient (r_Both_) and the linear regression line (dotted line) was made. The diagonal is indicated (grey line).

### Analytical robustness of ctDNA detection using UMIseq

To evaluate the analytical robustness of UMIseq, we assessed ctDNA detection in three settings using paired samples (**Figure 4E-G**). First, we tested the performance of UMIseq against an orthologous method – single mutation digital droplet PCR (ddPCR) – using two plasma aliquots (n = 11) from the sample blood collection. We found that 91% (10/11) of samples analyzed with UMIseq were ctDNA positive, while 64% (7/11) were ctDNA positive by ddPCR. The estimated cAFs of samples classified as ctDNA positive by both methods demonstrated a near-perfect correlation (r = 0.9950, Pearson’s correlation) (**Figure 4E**). Secondly, we tested the reproducibility of the entire workflow (from cfDNA purification through UMIseq calling) by procuring two plasma aliquots from 19 randomly chosen patients and repeating the entire workflow from cfDNA extraction to UMIseq call. For 84% (16/19) of the sample pairs, the UMIseq ctDNA classifications agreed. Additionally, a strong correlation was observed between the estimated cAFs in pairs where both samples were ctDNA positive (r = 0.9952, Pearson’s correlation) (**Figure 4F**). Lastly, we tested the repeatability of UMIseq, by splitting 49 randomly chosen cfDNA samples in two aliquots with identical amount of cfDNA which were then subjected to UMIseq analysis. Similar to the reproducibility analysis, we observed consistent ctDNA classification in 84% (41/49) and a strong correlation between the cAFs of the split samples (r = 0.9882, Pearson’s correlation) (**Figure 4F**).

### Validation of UMIseq

Having determined the UMIseq score threshold yielding 95% specificity for ctDNA detection in the training cohort, we next sought to validate whether this threshold would yield a similar specificity in an independent set of plasma from healthy individuals (n = 24). At the predefined UMIseq score threshold, the specificity estimates were indistinguishable from those of the training cohort (**Suppl. Figure 4** and **Suppl. Information**). We next conducted an independent validation of the UMIseq sensitivity on pre-OP plasma from stage I-III CRC patients (n = 209; stage I: 52, stage II: 108, stage III: 49). Overall, the sensitivity was of UMIseq was 70% (146/209) in the validation cohort. Stratifying the ctDNA detection rate based on UICC (**Figure 5A**), and pT and pN categories (**Figure 5B**) confirmed the association between the pT category and ctDNA detection (pT2: 56%; pT3: 75%; pT4: 81%) previously observed in the training cohort. To evaluate the potential for ctDNA detection in very low-stage cancers and premalignant disease, we applied UMIseq to pre-OP plasma obtained from patients diagnosed from fecal immunochemical test screening with minimally invasive CRC (pT1pN0) and colorectal adenomas. UMIseq detected ctDNA in the plasma of 31% (9/29) of the patients with pT1pN0 tumors but in none the patients with adenomas (**Figure 5C**).

**Figure 5.**
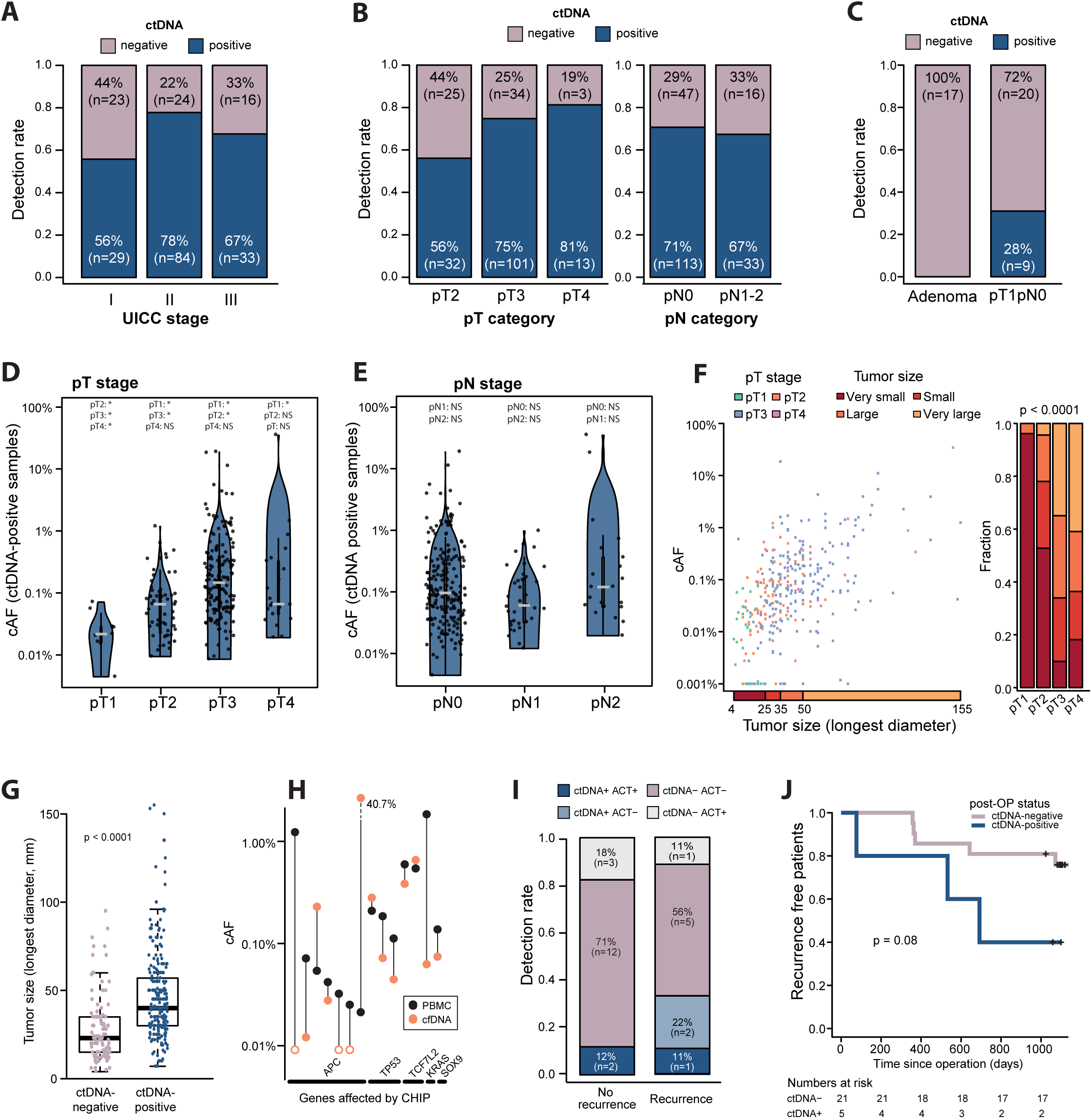
Validation cohort. **A)** ctDNA detection rates stratified by UICC stage. **B)** ctDNA detection rates stratified by pT stage (left), and pN stage (right). Note that one pT1N2M0 sample (ctDNA-negative) is omitted from the left panel. **C)** ctDNA detection rates in non-invasive colorectal adenomas and minimally invasive pT1N0M0 tumors. **D-E)** Estimated cAFs of all ctDNA-positive samples in the training and validation cohort (n = 223), stratified by pT **(D)** and pN **(E)** categories. Asterisks indicate significant differences (p < 0.05, Wilcoxon rank sum test) to other categories as annotated. Grey horizontal lines indicate median cAF. **F)** Left panel: correlation between tumor size and cAF. The analysis includes both ctDNA-positive and ctDNA-negative samples (n = 130). Twenty-six samples with undetected ctDNA signal (cAF = 0) are displayed with cAF = 0.001%. Right panel: The distribution among tumor sizes of each pT category, displayed as fractions. The quantiles of tumor sizes were used to designate tumors as very small, small, large, and very large, as indicated in left panel. The p-value from a Fisher’s exact test for the association between pT and size is shown. **G)** Tumor size stratified by ctDNA status. **H)** CHIP mutations were identified at 14 positions in 13 patients through UMIseq of PBMCs. cAF of the mutation in the PBMCs and in the cfDNA is shown. Open circles indicate cAF = 0. **I)** ctDNA detection rates in post-OP samples from recurrent and non-recurrent stage III CRC patients. Samples positive (ctDNA+) and negative (ctDNA-) for ctDNA are shown in grey and blue, respectively, and samples from patients that received adjuvant therapy (ACT+) indicated with darker colors. **J)** Kaplan-Meier plot of the recurrence free survival time after operation in patients with ctDNA-positive and ctDNA-negative post-OP samples. Patients were censored (indicated with crosses) after the last negative CT scan (30 months or more after operation).

### Association between pT, tumor size, and ctDNA

Pre-OP cfDNA samples from the cancer patients from the training and the validation cohorts (n = 364) were used to investigate associations between analytical and clinical factors and ctDNA. We found no association between the plasma sequencing depth or conversion efficiency and ctDNA detection. A trending association was observed between the number of tumor-specific mutations within the capture panel and ctDNA detection (p = 0.0967, OR 1.14).

Next, we investigated the ctDNA levels in relation to depth of invasion, spread to regional lymph nodes and tumor size. The estimated UMIseq cAFs of ctDNA-positive samples ranged from 0.004% to 71.9% (median 0.095%, IQR 0.048% - 0.319%). In general, the median cAF increased with the pT category, except for pT4, which only displayed an increased cAF compared to pT1 (median cAF: pT1 = 0.023% [IQR 0.013%]; pT2 = 0.070% [IQR 0.087%]; pT3 = 0.155% [IQR 0.331%]; pT4 = 0.070% [IQR 0.305%]) (**Figure 5D**). There was no significant difference between the median cAF and the pN category (median cAF: pN0 = 0.105% [IQR 0.296%]; pN1 = 0.064% [IQR 0.137%; pN2 = 0.105% [IQR 0.235%]) (**Figure 5E**). Furthermore, in a multivariable logistic model, pT but not pN were strongly associated with ctDNA detection with the odds ratio of detecting ctDNA being 3.41 in pT2 compared to pT1 (p < 0.001) and as much as 20.24 in pT4 (p < 0.001) (**Suppl. Table 3A**), an observation that complemented the pT-stratified and pN-stratified ctDNA detection rates (**Figure 4C**, **Figure 5B**). Even when adjusting for tumor size (longest diameter) a similar, but less strong, trend was observed (**Suppl. Table 3B**). In general, the cAFs increased with larger tumor size (**Figure 5F, left panel**). Furthermore, tumor size was strongly associated with the pT category, with more than 95% of pT1 tumors being classified as “very small tumors” while pT3 and pT4 tumors predominantly were “large tumors” and “very large tumors” (**Figure 5F, right panel**) (p < 0.0001, Fisher’s exact test). Lastly, the tumor size of ctDNA-negative patients (median 23.0 mm) was significantly smaller than the tumor size of ctDNA-positive patients (median 40.0 mm) (p < 0.0001, Wilcoxon rank sum test) (**Figure 5G**).

### Assessment of clonal hematopoiesis of indeterminate potential

As part of the UMIseq pipeline, potential mutations related to CHIP are identified in patient-matched normal PBMC DNA and excluded from the mutational catalogue and hence do not contribute to the UMIseq score of that patient. Since the PBMC DNA in this study was sequenced to the same depths as the plasma cfDNA (**Suppl. Figure 1C-D**), it allowed us to comprehensively evaluate the frequency of CHIP within the capture panel, i.e. in the most recurrently mutated regions in CRC. We found that 3.4% (13/381) of patients had CHIP mutations, comprising 14 distinct mutations in the genes *APC*, *TP53*, *TCF7L2*, *KRAS*, and *SOX9* (**Figure 5H**). In general, the cAF observed in PMBC DNA was higher than the cAFs observed in the cfDNA, with only four out of the 14 mutations having a higher cAF in cfDNAs. While the cAFs in PBMCs were generally low (less than 1%), *APC* was the only gene with cAFs less than 0.1%.

### UMIseq applied for minimal residual disease detection

We investigated the ability of UMIseq to detect MRD in post-OP plasma samples collected after operation and before initiation of adjuvant chemotherapy from patients with stage III CRC (n=26). The recurrence rate in this cohort was 31% (9/26). UMIseq detected ctDNA in 33% (3/9) of recurrence patients (**Figure 5I**) and post-OP ctDNA detection was associated with an increased risk of recurrence (p=0.08, log-rank test) (**Figure 5J**). ctDNA was detected in 12% (2/17) of non-recurrence patients, which translates to an apparent post-OP specificity of 89% (**Figure 5I**). However, both ctDNA-positive non-recurrence patients received adjuvant chemotherapy, which may have eliminated residual cancer cells left after the operation.

## DISCUSSION

Identification of cancer through ctDNA analysis has several clinical implications, and currently, the use of ctDNA analysis to detect and monitor residual disease after the operation is extensively investigated^3,5,18^. For successful clinical implementation in this setting, it is crucial for ctDNA analysis to exhibit analytical robustness and allow detection of low tumor burden (high sensitivity), while simultaneously minimizing the occurrence of false positives (high specificity). Additionally, the cost and simplicity of the analysis are inevitable factors that impact the utilization of ctDNA analysis. In this study, we introduced UMIseq. By using a fixed panel, designed to be applicable to all CRC patients, UMIseq provides a simple and time-efficient approach.

The key to sensitive ctDNA detection at a high specificity is a high signal-to-noise ratio. This can be achieved by increasing the signal, lowering the noise, or both. On one end, methods that aim at lowering the error rate, such as Duplex sequencing, achieve an error rate approaching 10^-7^. However, Duplex sequencing suffers from the inefficient recovery of both original strands, which occurs in a minority (usually 20–25%)^10,19^ of DNA fragments, thus limiting the sensitivity. On the other end, WGS approaches aim to increase the signal by integrating the genome-wide signal for ctDNA detection. Performance comparisons between the different approaches are challenging due to the lack of adequate benchmarking studies, leaving the comparisons to be made between studies and at the cohort level. This has multiple weaknesses *e.g.*, that plasma volumes, sample processing, and cohort compositions are different. The analytical sensitivities observed in plasma samples are, a bit surprisingly, quite similar for targeted and WGS approaches - often the lowest detected AFs are in the range 10^-4^ to 10^-5^, indicating that in practice both targeted sequencing and WGS strategies have their own limitations^11,12,14,20,21^. Presumably, targeted strategies that employ comprehensive error correction through UMIs or Duplex sequencing are limited by the amount of signal captured for ctDNA detection, while WGS approaches are limited by their error rate despite comprehensive error modeling and correction. Similar to other studies, UMIseq enabled ctDNA detection down to a cAF below 10^-4^ with the lowest observed cAF in a ctDNA positive sample being 0.004% and a theoretical LOD below 0.001% in 10% of the estimated LODs at 20,000x sequencing depth.

Similar to findings in other studies^13,21,22^, the error rate of UMIseq was highly dependent on nucleotide substitution and trinucleotide context. Particularly, C>T substitutions in the context of N(C>T)G were error-prone, while T>G and C>G substitutions, regardless of trinucleotide context, had a low error rate. In general, INDELs, especially insertions, had a lower error rate than SNVs, which was also reflected in the theoretical LOD calculation. Although these observations are based on a fixed hybrid-based capture panel, similar trends have been observed from ddPCR^22^ and are expected to be generalizable to other approaches as well. This suggests that a preference towards low error rate substitutions, with regard to the trinucleotide context, and the inclusion of non-SNV targets could further enhance the sensitivity of ctDNA detection.

A potential challenge to maintaining the specificity of ctDNA testing is the risk of false positive ctDNA calls by misinterpreting CHIP as ctDNA mutations^23–25^. Studies employing deep sequencing of PBMCs from healthy individuals suggest that low-frequency CHIP mutations (<0.1%) are present in up to 92% of patients^23,25^. When utilizing a tumor-informed strategy for ctDNA detection, only CHIP-related mutations that overlap with tumor-specific variants can contribute to false positive ctDNA calls. From tumor-informed UMIseq analysis of PMBC DNA, we identified and excluded CHIP-related mutations in 3.4% (13/381) of the patients. These results demonstrate that while CHIP-related mutations can still be an issue for a tumor-informed strategy, it is far less likely to result in a false positive call than when using a tumor-agnostic approach. To minimize the additional cost and labor associated with CHIP analysis, it may be feasible to restrict CHIP analysis to patients with a ctDNA-positive sample.

Previous studies have demonstrated that the ctDNA detection rate increases with the UICC stage and tumor size^26,27^. While our study supports these observations, we find that this correlation is primarily driven by the pathological T stage. This is corroborated by the lower detection rate of pT1pN0 tumors and the absence of ctDNA in patients with non-invasive colorectal adenomas. In both the training and validation of UMIseq, we surprisingly found a complete lack of association between ctDNA detection and lymph node tumor infiltration (pN stage), although we cannot rule out that a small association would be identified in a larger patient population. Similar observations have been noted in gastric cancers^28^. One interpretation of this is that the additional tumor burden from regional lymph node involvement does not contribute to increased blood ctDNA levels, suggesting that tumor cells residing in the lymph nodes shed very limited amounts of cfDNA into the bloodstream. Possibly, this can be attributed to an abundance of immune cells in the lymph nodes, which degrade cell residues, including any ctDNA released by the tumor cells. A potential consequence could be a reduced ability to detect residual tumor cells residing in lymph nodes after operations, which is consistent with recent observation in a study of colorectal metastases^29^.

We developed UMIseq with the aim of providing a sensitive and uniform approach for detection and surveillance of residual disease in CRC patients after the operation. In addition to thoroughly evaluating the performance of UMIseq, we investigated the potential for MRD detection in post-OP plasma samples. This revealed a sensitivity of 33% and a specificity of 89% for post-OP ctDNA detection. While UMIseq did enable detection of MRD already immediately after the operation, the ctDNA detection at this timepoint is possibly challenged by a very low tumor burden and a surge in release of trauma-related cfDNA^30^. Through serial surveillance, ideally using plasma samples collected after end of adjuvant chemotherapy, the sensitivity is expected to improve. The lower specificity, compared to the specificity observed in the training and validation cohort, is expected in stage III CRC as this patient group usually receives adjuvant chemotherapy after the collection of the post-OP sample^3,31,32^, which was also the case for the two ctDNA-positive non-recurrence patients. Here, we explored MRD detection in a small cohort of stage III CRC patients, and further studies are required to assess the clinical utility of UMIseq in the post-operative setting across all stages.

There are limitations to this study. First, the targeted panel applied for UMIseq was specifically designed for CRC patients and can therefore not be applied to other cancer types in its current design. However, by designing a new capture panel, UMIseq could be easily adapted for other cancer types while keeping the protocol for sample preparation and the bioinformatic pipeline consistent. The required size of such a panel should be altered according to cancer type and may result in an increase in sequencing cost. The panel designed for this study was tailored for identification of small somatic mutations. As off-target persists as a relatively large fraction (∼15%) within the sequencing data, there is potential to leverage this information and gain insight into patient-specific copy number alterations. This could provide a more comprehensive patient-specific tumor catalogue for improved ctDNA detection. While we demonstrate the overall robustness of UMIseq, the reproducibility and repeatability assessment were conducted within a single laboratory, preventing evaluations of the inter-laboratory robustness of UMIseq.

In summary, we here present UMIseq as a sensitive and universal approach for ctDNA detection, where the same laboratory workflow and capture panel are applied to all CRC patients. In this study, we thoroughly assessed the performance of UMIseq using pre-operative plasma samples and investigated the potential of UMIseq for MRD detection. The potential application of this approach is aimed at detecting MRD and monitoring for recurrence, which is not limited to CRC but also in other cancers, provided an appropriate capture panel for the given cancer type.

## ADDITIONAL INFORMATION

## Acknowledgments

We thank the blood donors, CRC patients, and the Danish Cancer Biobank for contributing clinical material. We also thank the IMPROVE study group for patient inclusion: Kåre Andersson Gotschalck, Lene Hjerrild Iversen, Uffe Schou Løve, Claudia Jaensch, Ole Thorlacius-Ussing, Per Vadgaard Andersen.

## Author contributions

AF, MHR, IT, and CLA designed the study. SØJ contributed to patient selection and planning. AF and MHR were responsible for sample preparation and sequencing. MHR developed the bioinformatic pipeline for UMIseq analysis. TVH and CD contributed to data acquisition and interpretation. HD, HH, and LD contributed to patient recruitment. MD and JSP contributed to data interpretation. AF and MHR analyzed the data and drafted the manuscript, which was revised by all authors.

## Ethics approval and consent to participate

The study was approved by the Danish National Committee on Health Research Ethics (case no. 2208092). The study was performed in accordance with the Declaration of Helsinki.

## Consent for publication

All participants provided written informed consent.

## Data availability

The sequencing data generated during the study will be available through controlled access from GenomeDK (https://genome.au.dk/library). All other data supporting the findings of this study are available upon reasonable request to the corresponding author.

## SUPPLEMENTARY FIGURES

**Suppl. Figure 1.**
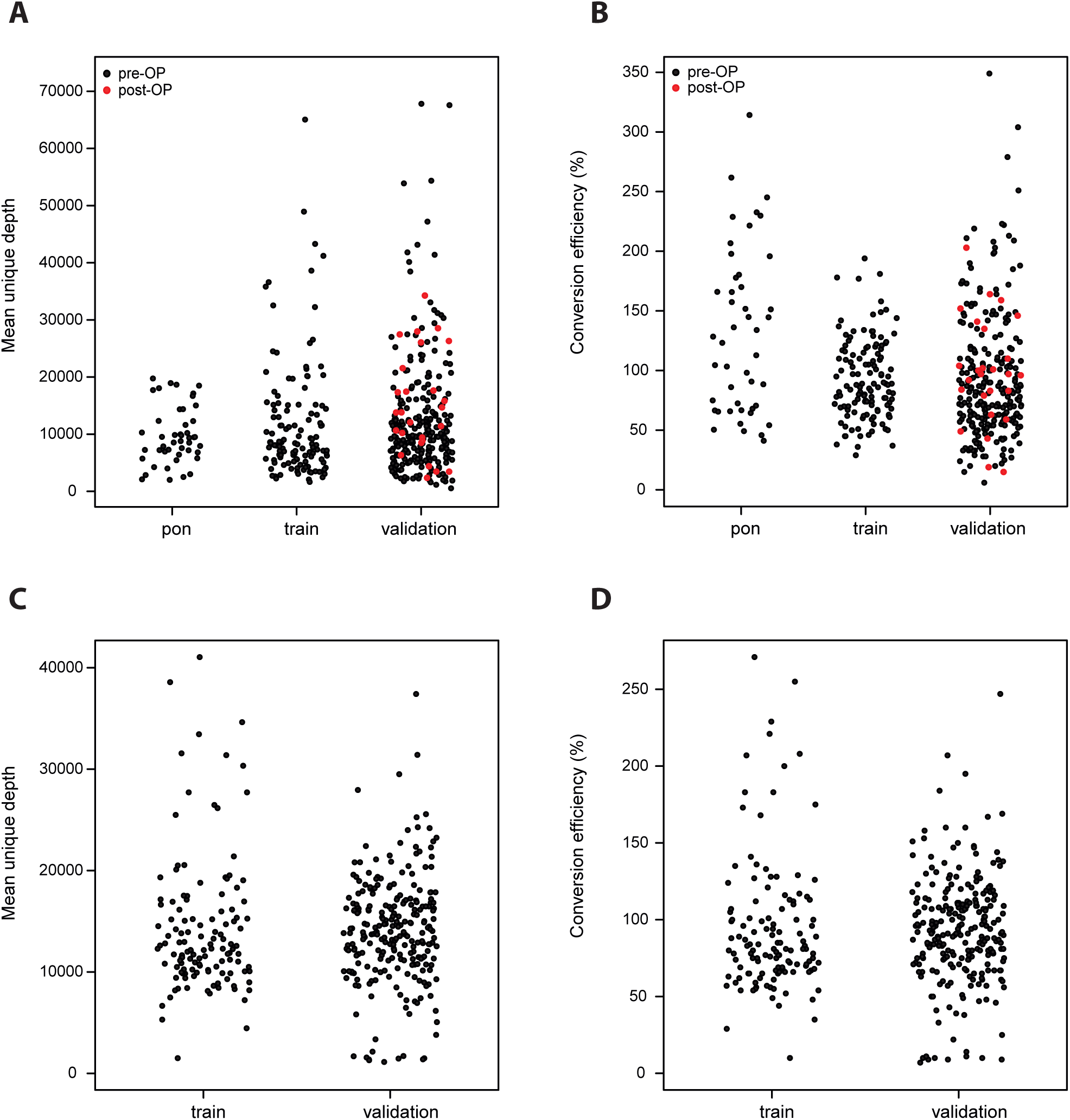
Coverage and conversion efficiency of plasma and PBMC samples. **A)** Mean coverage of plasma samples in the PON, training and validation cohorts after UMI consensus generation. **B)** Conversion efficiency (*i.e.* the mean consensus sequencing depth divided with genome equivalent used as input to library) of plasma samples in the PON, training and validation cohorts. The theoretical maximum conversion is 200%, which corresponds to where the Watson and Crick strands of all double stranded molecules of the input cfDNA have been sampled and converted into sequence data (See **Supplementary Information**). In **A-B)** the post-OP samples are indicated in red. **C)** Coverage of PBMC samples in the training and validation cohort after UMI consensus generation. **D)** Conversion efficiency of PBMC samples in the training and validation cohort.

**Suppl. Figure 2.**
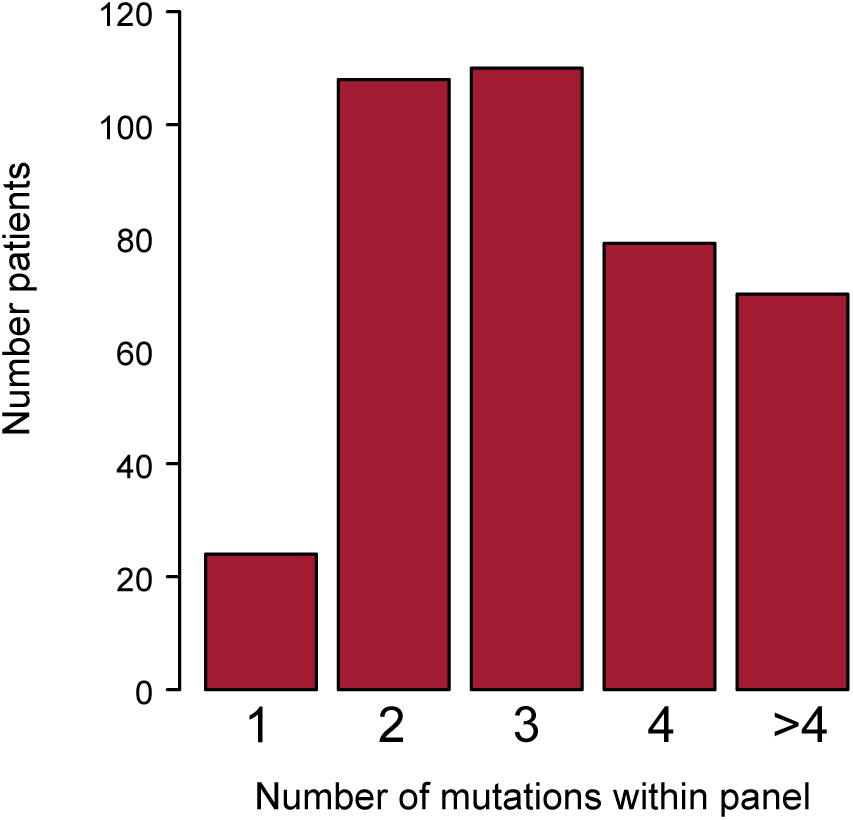
The distribution of SNV and non-SNV mutations. The distribution of SNVs (n) and non-SNVs (k) on the panel per patient in the study (n = 381). The individual numbers show the frequency of SNV and non-SNV combinations, as also reflected by the color scale.

**Suppl. Figure 3.**
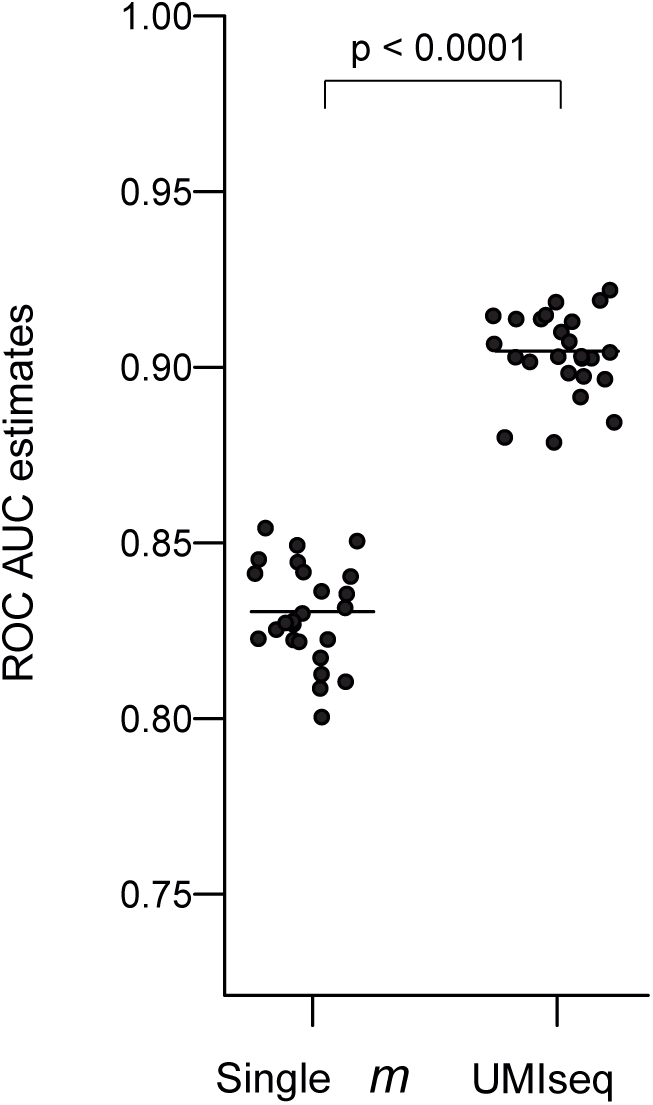
Mutations per patient. Number of tumor-specific mutations (SNV, INDELs, MNVs) within the capture panel for each patient of the study (n = 381).

**Suppl. Figure 4.**
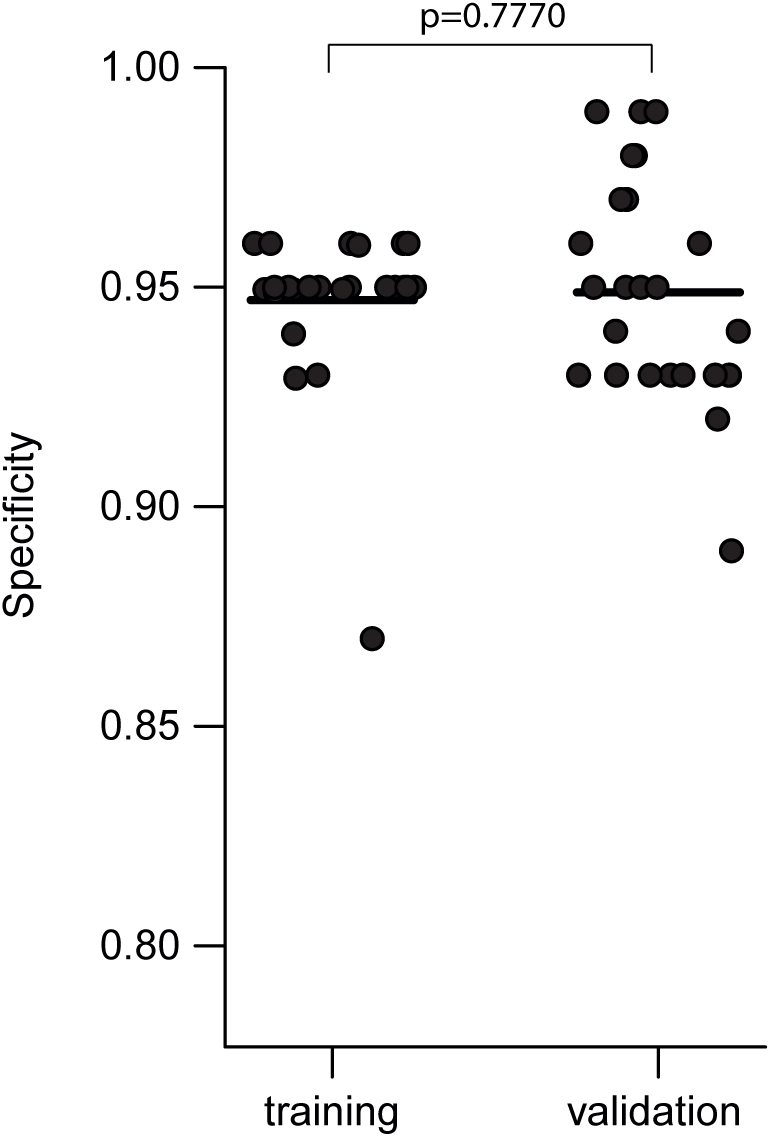
ROC AUC estimates for single mutation and UMIseq strategies. ROC AUC estimates for UMIseq were estimated by Monte Carlo simulations (N = 25) using the healthy controls (n = 37) and CRCs (n = 126) samples of the training cohort as described in Materials and Methods. To estimate the performance of a strategy that only uses a single mutation marker (*‘*single *m*’), we used the mutation with the lowest *m* score (strongest mutational signal) in each plasma sample (both cancer and non-cancer control samples) directly as the sample score. The ROC AUC from each simulation was finally calculated. A student’s t-test was applied to test the difference in mean ROC AUC for the single *m* and UMIseq strategy.

**Suppl. Figure 5.**
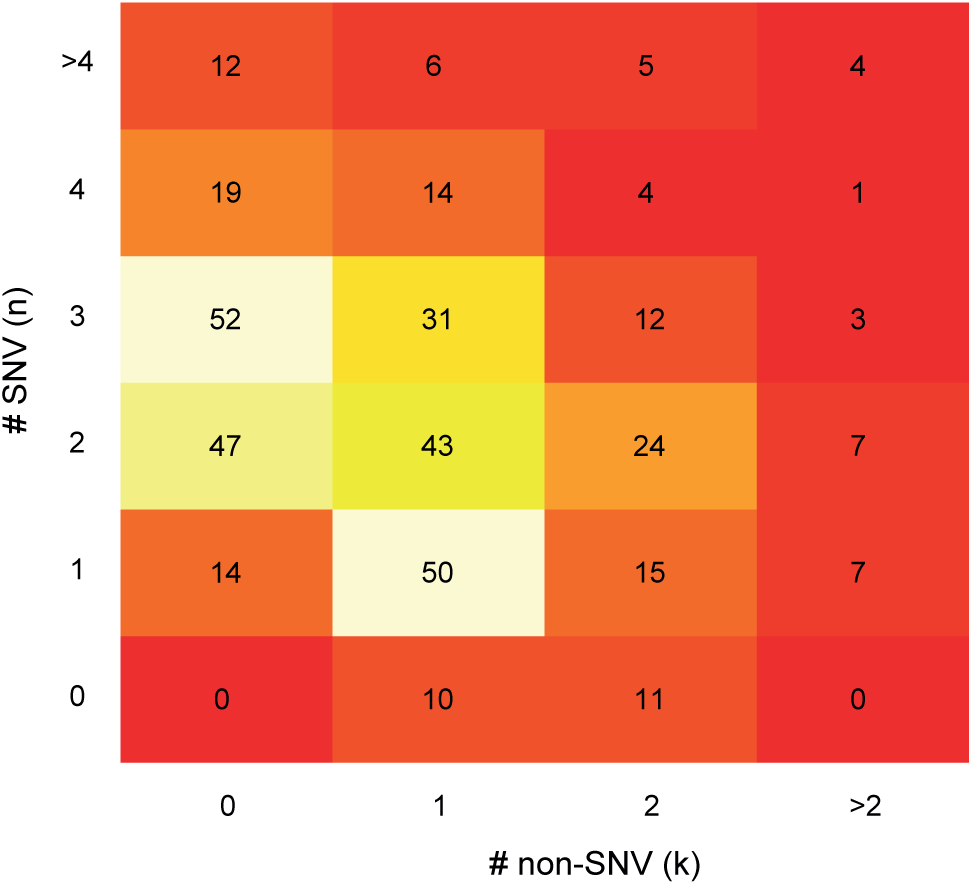
UMIseq specificity of the training and validation cohorts. Each point is the UMIseq specificity estimate from a Monte Carlo simulation (N = 25 in each cohort) in the healthy controls of the training (n = 37) and validation (n = 24) cohorts. For each simulation, n = 100 UMIseq scores S was calculated from random negative sample sets generated by sampling mutations from the training tumor catalog (n = 276 unique mutations) using the same procedure as described for the training of the UMIseq algorithm (see Materials and Methods). The specificity was calculated as the fraction of the 100 *S*-scores below the UMIseq threshold (*i.e.* resulted in a negative call). A student’s t-test was applied to test for a difference in the mean of the training and validation specificity.

## SUPPLEMENTARY INFORMATION

### Tumor and normal DNA sequencing library preparation

DNA was extracted from formalin-fixed paraffin-embedded (FFPE) tissue using QiAamp DNA FFPE tissue kit (Qiagen), from fresh frozen (FrFr) tissue using Puregene DNA Purification Kit (Gentra Systems), and from PBMC (normal DNA) using the QIAamp DNA Blood Kit (Qiagen). Tumor and normal DNA were quantified using the Qubit™ dsDNA BR Assay Kit (ThermoFisher).

Tumor and normal DNA sequencing libraries were generated using xGen UDI-UMI Adapters (Integrated DNA Technologies Inc., IDT) and the Twist Library Preparation Enzymatic Fragmentation Kit 1.0 (TWIST Bioscience). Libraries were prepared as described by the manufacturer. For normal and FrFr DNA, we used 50 ng input and 10 min fragmentation. For FFPE DNA, 200 ng input and 6 min fragmentation were used. All libraries were amplified with seven cycles of PCR. Libraries were quantified using Qubit™ dsDNA BR Assay Kit (ThermoFisher), and library size was estimated using TapeStation D1000 (Agilent).

### Tumor and normal DNA sequencing and variant calling

Tumor and matched normal DNA were whole-exome sequenced (WES) as follows. Libraries were captured using the NGS Human Core Exome (TWIST Bioscience, ∼33 Mb) according to the manufacturer’s protocol. Target-enriched libraries were sequenced using the NovaSeq platform with 2×150 bp paired-end sequencing to a target depth of 150x for FrFr tissue DNA, 200x for FFPE tissue DNA, and 80x for normal DNA. Raw sequencing reads were converted into FastQ files using Illumina bcl2fastq. Sequencing adapters were removed bioinformatically by cutadapt (v3.0)^1^, and trimmed reads were mapped to the human reference genome (hg38) using BWA-MEM (v0.7.17)^2^ with PCR duplicates flagged by Picard (v2.23.3) MarkDuplicates^3^. Alignments were processed further using GATK (v4.1.9.0) BaseRecalibrator according to the GATK Best Practices. Patient-specific single nucleotide polymorphism (SNPs) were identified with GATK HaplotypeCaller^4^, scored with GATK CNNScoreVariants^5^, and filtered by GATK FilterVariantTranches^6^. Somatic SNVs and small INDELs were identified using GATK Mutect2^7^ and Strelka2^8^.

For patients with colorectal adenomas (n = 17), tumor and normal DNA were whole-genome sequenced (WGS) to a target depth of 30x and 20x, respectively. The bioinformatic processing was the same as described above.

For a subset of the cancer patients (n = 19), the tumor and matched normal DNA were sequenced with targeted duplex sequencing as previously described by Christensen et al.^9^, using the same capture panel as for plasma cfDNA (see below). Duplex sequencing data were analyzed using the same bioinformatic pipeline used for plasma cfDNA (see below), with the exception that for assessing variants the stringent Shearwater ‘OR’ model was used, and that mutations were called across the whole panel (i.e. tumor agnostic). A somatic tumor mutation was called when the following three conditions were fulfilled: (1) the tumor DNA *m*-score (see below) was below or equal to 0.01, (2) the normal DNA *m*-score was above 0.01, and (3) the normal DNA allele frequency (AF) was below 5%.

### Sample collection for UMIseq

Blood samples from patients and healthy individuals were collected in 10 mL K2-EDTA tubes (Becton Dickinson). Plasma was isolated within 2 hours of blood collection by double centrifugation at 4000 rpm for 10 min and stored at −80 °C until DNA extraction. Plasma cfDNA was extracted using the QiaAMP Circulating DNA kit (Qiagen) and quantified using digital PCR as previously described^10^. Prior to cfDNA extraction, all plasma samples were spiked with a fixed number of soybean CPP1 DNA fragments, which was used to estimate purification efficiency, as previously described^11^. The cfDNA fragment size distribution was assessed using cfDNA ScreenTape (Agilent).

### UMIseq library preparation

UMIseq sequencing libraries were generated for both plasma cfDNA and PBMC DNA (used for filtering clonal hematopoiesis of indeterminate potential [CHIP] mutations) from all patients and healthy individuals. Normal DNA libraries were prepared as described above for WES.

Plasma cfDNA libraries were made using cfDNA from 8 mL of plasma, with an upper cap of 80 ng cfDNA per library and a maximum of two libraries per plasma sample. Libraries were generated using xGen UDI-UMI Adapters (IDT) and KAPA HyperPrep kit (Roche). Post-ligation clean-up was performed with AMPURE beads in a 1.4x (beads/DNA ratio) to retain short fragments, while post-PCR clean-up was done using a 1.0x ratio. The libraries were split into four separate reactions during PCR and amplified with seven cycles of PCR. Libraries were quantified using Qubit™ dsDNA BR Assay Kit (ThermoFisher) and library fragments size was estimated using TapeStation D1000 (Agilent).

### Targeted capture of UMIseq libraries

Sequencing libraries generated from cfDNA and PBMC DNA were subjected to targeted capture. We employed a hybrid-based capture approach using a custom panel (Roche CapSeq custom panel, Roche), targeting the most frequently mutated genomic regions observed in patients with CRC^12^, and designed to capture at least one mutation in 99% of CRC patients. The capture panel covered 42 exonic regions in 12 genes (15,465 bp). Additionally, for sample tracking purposes the panel incorporated 18 single-nucleotide polymorphisms (SNPs) (2,629 bp) with a high heterozygosity index in the general Danish population (**Suppl. Table 1**). Throughout the manuscript, we will solely refer to the cancer-centric part of the panel.

The capture reaction was carried out according to the manufacturer’s protocol, with the following modifications. To create each capture pool, we combined 4-12 libraries utilizing 12 µL of each library (accounting for ∼25% of the total library volume). We ensured that each capture pool contained a maximum of 5000 ng library. Two consecutive capture reactions were performed by repeating the entire capture protocol, using half of the probe volume for each capture reaction. This approach was employed to maximize the on-target fraction (>85%), as a single capture alone did not achieve the desired level of efficiency (< 25%). The captured libraries were quantified and size estimated using TapeStation HS.

### Sequencing and processing of UMIseq libraries

The captured libraries underwent 2×150bp paired-end sequencing using the Illumina NovaSeq platform. To ensure comprehensive sequencing coverage, all libraries were sequenced to saturation, meaning that the entire complexity of the captured library was sequenced, and each unique fragment was sequenced multiple times. This strategy facilitated the generation of UMI families comprising of at least 3 reads for >99% of data.

To determine the amount of data (in gigabases [Gb]) required to achieve sufficient saturation for each sample, we utilized the following formula: Gb = *N* * *P* * *Fs* * *d*. Here, *N* represents the genomic equivalents (GEs) used as input for the library, *P* denotes the size of the capture panel, *Fs* represents the desired median UMI family size (which was set at 12 reads), and *d* is an empirically determined factor, which was set at 3 and ensures sufficient data for all samples in the capture pool. *P*, *Fs*, and *d* remain constant across samples, while *N* varies for each library. To determine the Gb necessary for saturating all samples in a capture pool, we identified the sample within the pool that required the most data and multiplied that by the number of samples in the pool. Although this approach resulted in a higher *Fs* value than strictly necessary, it guaranteed that all samples received adequate data and obviated the need for re-sequencing.

Fastq reads were mapped to hg38 using BWA-MEM (version 0.7.17)^2^. Next, reads with similar UMI and genomic mapping position were grouped together using the umitool group (version 1.1.2)^13^ with the directional method and edit-distance-threshold=1, before collapsing individual UMI family reads into consensus reads with fgbio (version 2.1.0) commands CallMolecularConsensusReads (parameters: --error-rate-pre-umi=45, --error-rate-post-umi=40, and --min-input-base-quality=10) and FilterConsensusReads (parameters: --min-reads=3, --max-base-error-rate=0.1, --max-read-error-rate=0.01, --min-base-quality=15, --min-mean-base-quality=20 and --max-no-call-fraction=0.2). The consensus reads were remapped to hg38 using BWA-MEM and realigned with Abra2 (version 2.24)^14^. Finally, overlapping read pairs were clipped using fgbio ClipBam^15^, to create the UMI consensus BAM file. The forward and backward mutation and non-mutational read counts that intersected with the patient-matched tumor VCF file were extracted from the consensus BAM file using a custom script.

### Conversion efficiency of cfDNA libraries

To assess how much of the original cfDNA, extracted from 8 mL of plasma, was converted into useful UMI collapsed reads, the conversion efficiency was calculated as conversion = consensus_sequencing_depth / input_ge, where consensus_sequencing_depth is the mean UMI consensus sequencing depth obtained for a sample, and input_ge is the number of genome equivalents that was used as input in the cfDNA library. Since UMIseq utilizes NGS adapters with single stranded UMIs (on the P7 adapter arm), the theoretical maximum conversion is 200%, which corresponds to a situation where the Watson and Crick strands of all double stranded molecules of the input cfDNA have been sampled and converted into sequence data.

### Sample concordance check

Each set of patient samples (tumor, normal, cfDNA) was evaluated to confirm common patient origin. This was done by assessment of 18 SNPs (**Suppl. Table 1**). For each patient, the SNP genotype of the normal sample was compared to the SNP genotype of the tumor and cfDNA samples. No mismatches were observed.

### Blacklisting of SNVs and INDELs

Out of 46,395 possible SNVs (capture panel size times possible base changes = 15,465 bp * 3), we blacklisted 69 substitutions (0.15%) because they had a cAF above 1% in more than one PON sample, i.e., they were recurrently noisy in plasma cfDNA from healthy individuals. In addition, we blacklisted 45 SNPs, and 24 regions affected by frequent insertions or deletions in the plasma cfDNA from healthy individuals. Blacklisted SNVs, SNPs, and INDELs were excluded when analyzing patient plasma samples.

### Flagging of mutations associated with clonal hematopoiesis

For mutations associated with clonal hematopoiesis, it is not possible to determine if a signal seen in the matching cfDNA, has tumor and/or hematopoietic origin. Therefore, potential CHIP mutations were excluded from further analysis by the following approach.

Variants with a mutation score *m* below 0.05 in patient-matched PBMC samples, were flagged as potentially representing CHIP. In total, 14 variants (representing 14 distinct mutations in 13 patients) overlapped the paired tumor sample mutation catalogs. These were marked as CHIP and excluded from further analysis. In addition, one mutation (chr10:113165610_C/A, TCF7L2_Pro483Thr) was flagged as CHIP in two control PBMC samples and were excluded from the corresponding cfDNA control data sets.

### Limit of detection calculation of individual mutations

The limit of detection (LOD) of a particular mutation was defined as the minimum number of mutational reads required for the mutation to be called at a total depth of 20,000 with a false positive error rate of 1% against the PON. To explore the difference in LODs for SNVs and single-nucleotide INDELs, the mutational score *m* corresponding to 1% false positive rate (FPR) was first identified for SNV and INDELs separately. Then, the LOD for 1,000 random SNV and INDEL mutations was calculated. A two-sample Kolmogorov-Smirnov test was used to test the similarity of the underlying LOD distributions. In addition, the LODs for selected CRC-related mutations with a recurrence index above 1% in the Catalogue of Somatic Mutations in Cancer (COSMIC) were calculated and added for comparison.

### In silico estimation of the ctDNA detection probabilities in mutation catalogs using SNVs or both SNVs and non-SNVs

For each patient in the study, the number of SNVs (n) and non-SNVs (k; INDELs and MNVs) mutations occurring within the panel were counted to make an n,k-distribution (**Suppl. Figure 5**). Twenty-five *in silico* mutation catalogs were generated by drawing random SNV and non-SNV mutations from the combined mutation catalog of the patients in the study. The number of SNVs and non-SNVs in each catalog was determined by sampling in the n,k-distribution. As the analysis required the presence of both SNV and non-SNV mutations, catalogs where either n or k equaled zero were discarded. Next, the mutation catalogs were populated with *in silico* counts of forward/backward alternative counts (ALT/alt) and normal counts (^ALT/^alt) by sampling from a Poisson distribution with λ = cAF (alternative counts) or λ = 1-cAF (normal counts) to reach a depth of 20,000x. From each of the total 25 mutation catalogs, two integrated *s*-scores were generated: one using the mutation catalog (including non-SNV mutations) and one including only the SNV mutations, thus generating 50 scores. Five different cutoffs were then applied to call the catalogs positive or negative at increasing specificities. This cutoff was arbitrarily set at the 5, 25, 50, 75, and 95 percentiles of all the 50 generated scores. Finally, a logistic regression line was fitted to the calls of the SNV plus non-SNV, and SNV only sets in order to visualize the probabilities as a function of specificity. This procedure was repeated with four different cAFs: 0.1667%, 0.0556%, 0.0185%, and 0.0062%.

### Recurrent COSMIC mutations

Mutations in COSMIC(12) (release v98) were retrieved from cancer.sanger.ac.uk. Records were filtered for being from a WES or WGS effort (’Genome.wide.screen’); ‘Primary Site’ equal "large_intestine"; ‘Sample.Type’ being either of "surgery fresh/frozen", "NS", "surgery-fixed"; having a Pubmed id (’Pubmed_PMID’); from a large study (more than 10000 mutation records); the mutation (’Mutation.Description’) being either of "Substitution - Missense" or "Substitution - Nonsense". Records were translated to the genomic positions of the sense strand and intersected with the capture panel. The recurrence rate of individual records was calculated as their frequency divided by the total number of samples (’ID_sample’), and recurrent mutations were defined as mutations with more than one occurrence in the data set.

### Analytical sensitivity analysis using a synthetic mixture

Tumor and matched PBMC DNA from 36 patients were Duplex sequenced, as previously described(9), to a median mean depth of 2284x (range 113x - 7323x) using the UMIseq capture panel and data processed and calls generated as described above for tumor and normal DNA sequencing. To create a compound sample (Sample A) with a foreground set of ground truth mutations, DNA from all 36 tumors was mixed in equal volumes. Sample A was serially diluted with normal PBMC DNA from a healthy donor (Sample X, diluent). The serial dilution of Sample A with Sample X was done in 1:3 ratios to generate six samples with decreasing mutational signals (Sample A through Sample F, **Figure 3A**). UMIseq libraries of the samples A-F and X were generated using a DNA input of 25,000 GEs. The libraries were sequenced to a median depth of 16,638x (range: 15,758x - 23,864x) and variants were called using Shearwater. The foreground set of mutations was defined by first identifying variants (SNPs and SNVs) with a total sequence depth above 30x and an AF above or equal to 1% in each of the 36 tumors. Further, variants that were supported by less than 10 reads in Sample A, as well as variants identified in Sample X, were blacklisted. This resulted in 95 high-confidence mutations of sample A, which were used as a foreground. All other possible substitutions within the panel, except blacklisted variants, were used as background. The foreground mutations were grouped based on their observed AF across sample A through F. These subgroups comprised six AF ranges: 0-0.01%, 0.01%-0.05%, 0.05-0.1%, 0.1-0.5%, 0.5-1%, and 1-100%. (**Figure 3A**). The number of mutations in each interval was: 32, 86, 58, 90, 40, and 52. Precision-recall and receiver operator characteristic curve estimates were generated for every interval by 10 Monte-Carlo simulations with foreground mutations as positive labels, and random background mutations as negative labels, keeping the number of negative labels similar to the number of positive labels.

### Limit of detection calculation for UMIseq

The sample level UMIseq LOD was defined as the minimal cAF for a given patient mutation catalog that results in a UMIseq score which renders the sample ctDNA positive (i.e. above the UMIseq threshold). A distribution of LODs for each sequencing depth (5,000x, 10,000x, 20,000x and 40,000x) was estimated from n=100 *in silico* generated mutation catalogs (i.e. simulated samples). Each were made by selecting mutations so that both the number as well as the panel index of the mutations reflected the frequencies observed in the combined mutation catalog of the patients in the study (**Suppl. Figure 5**). Clonal mutations are, in general, expected to be associated with a higher frequency in plasma as compared to sub-clonal mutations. We simulated this effect by normalizing the tumor AFs to the highest AF among each tumor to generate a clonality factor in the (0,1]-interval where higher values indicate a more clonal mutation. For recurrent mutations the arithmetic mean across samples harboring that mutation was used. For each *in silico* mutation catalog, the minimum number of mutation counts - scaled according to the clonality factor - that exceeded the UMIseq threshold were identified. The empirical **S^N^**-distribution used to generate the UMIseq score was randomly chosen among all CRC plasma samples of the study for every *in silico* catalog. Finally, the corresponding cAF was calculated as described, and reported as the LOD for the particular *in silico* mutation catalog.

### Assessment of UMIseq robustness

#### Orthologous method test

From 11 patients, cfDNA was extracted from an additional 8 mL plasma aliquot and analyzed with single-marker ddPCR. The ddPCR approach, including assay design, cycling optimization, and error modeling was performed as previously described^16,17^. For the comparison of UMIseq and ddPCR, the cAF for ddPCR was reported for the single assay mutation, whereas the cAF for UMIseq was calculated as described above.

#### Reproducibility test

For a subset of blood samples (n = 19), cfDNA was extracted from a separate 8 mL plasma sample and analyzed with UMIseq. Sample calls and cAF levels of these paired plasma samples were used to evaluate the reproducibility of the UMIseq protocol.

#### Repeatability test

For 46 plasma samples, the cfDNA was split into two libraries with identical cfDNA inputs, which were then processed in parallel. Sample calls and cAF levels between these paired libraries were used to assess the analytical repeatability of UMIseq.

## REFERENCES

1 Alix-Panabières, C. & Pantel, K. Liquid Biopsy: From Discovery to Clinical Application. Cancer Discov 11, 858–873, doi:10.1158/2159-8290.Cd-20-1311 (2021).

2 Hasenleithner, S. O. & Speicher, M. R. A clinician’s handbook for using ctDNA throughout the patient journey. Mol Cancer 21, 81, doi:10.1186/s12943-022-01551-7 (2022).

3 Henriksen, T. V. et al. Circulating Tumor DNA in Stage III Colorectal Cancer, beyond Minimal Residual Disease Detection, toward Assessment of Adjuvant Therapy Efficacy and Clinical Behavior of Recurrences. Clin Cancer Res 28, 507–517, doi:10.1158/1078-0432.Ccr-21-2404 (2022).

4 Gale, D. et al. Residual ctDNA after treatment predicts early relapse in patients with early-stage non-small cell lung cancer. Ann Oncol 33, 500–510, doi:10.1016/j.annonc.2022.02.007 (2022).

5 Kotani, D. et al. Molecular residual disease and efficacy of adjuvant chemotherapy in patients with colorectal cancer. Nat Med 29, 127–134, doi:10.1038/s41591-022-02115-4 (2023).

6 Tarazona, N. et al. Targeted next-generation sequencing of circulating-tumor DNA for tracking minimal residual disease in localized colon cancer. Ann Oncol 30, 1804–1812, doi:10.1093/annonc/mdz390 (2019).

7 Phallen, J. et al. Direct detection of early-stage cancers using circulating tumor DNA. Sci Transl Med 9, doi:10.1126/scitranslmed.aan2415 (2017).

8 Ryoo, S. B. et al. Personalised circulating tumour DNA assay with large-scale mutation coverage for sensitive minimal residual disease detection in colorectal cancer. Br J Cancer, doi:10.1038/s41416-023-02300-3 (2023).

9 Kinde, I., Wu, J., Papadopoulos, N., Kinzler, K. W. & Vogelstein, B. Detection and quantification of rare mutations with massively parallel sequencing. Proc Natl Acad Sci U S A 108, 9530–9535, doi:10.1073/pnas.1105422108 (2011).

10 Newman, A. M. et al. Integrated digital error suppression for improved detection of circulating tumor DNA. Nat Biotechnol 34, 547–555, doi:10.1038/nbt.3520 (2016).

11 Newman, A. M. et al. An ultrasensitive method for quantitating circulating tumor DNA with broad patient coverage. Nat Med 20, 548–554, doi:10.1038/nm.3519 (2014).

12 Wan, J. C. M. et al. ctDNA monitoring using patient-specific sequencing and integration of variant reads. Science Translational Medicine 12, eaaz8084, doi:doi:10.1126/scitranslmed.aaz8084 (2020).

13 Christensen, M. H. et al. DREAMS: deep read-level error model for sequencing data applied to low-frequency variant calling and circulating tumor DNA detection. Genome Biol 24, 99, doi:10.1186/s13059-023-02920-1 (2023).

14 Kurtz, D. M. et al. Enhanced detection of minimal residual disease by targeted sequencing of phased variants in circulating tumor DNA. Nat Biotechnol 39, 1537–1547, doi:10.1038/s41587-021-00981-w (2021).

15 Forbes, S. A. et al. COSMIC: somatic cancer genetics at high-resolution. Nucleic Acids Res 45, D777–d783, doi:10.1093/nar/gkw1121 (2017).

16 Gerstung, M., Papaemmanuil, E. & Campbell, P. J. Subclonal variant calling with multiple samples and prior knowledge. Bioinformatics 30, 1198–1204, doi:10.1093/bioinformatics/btt750 (2014).

17 Gerstung, M. et al. Reliable detection of subclonal single-nucleotide variants in tumour cell populations. Nat Commun 3, 811, doi:10.1038/ncomms1814 (2012).

18 Tie, J. et al. Circulating Tumor DNA Analysis Guiding Adjuvant Therapy in Stage II Colon Cancer. N Engl J Med 386, 2261–2272, doi:10.1056/NEJMoa2200075 (2022).

19 Schmitt, M. W. et al. Detection of ultra-rare mutations by next-generation sequencing. Proc Natl Acad Sci U S A 109, 14508–14513, doi:10.1073/pnas.1208715109 (2012).

20 Zviran, A. et al. Genome-wide cell-free DNA mutational integration enables ultra-sensitive cancer monitoring. Nat Med 26, 1114–1124, doi:10.1038/s41591-020-0915-3 (2020).

21 Bae, J. H. et al. Single duplex DNA sequencing with CODEC detects mutations with high sensitivity. Nat Genet 55, 871–879, doi:10.1038/s41588-023-01376-0 (2023).

22 Henriksen, T. V. et al. Error Characterization and Statistical Modeling Improves Circulating Tumor DNA Detection by Droplet Digital PCR. Clin Chem 68, 657–667, doi:10.1093/clinchem/hvab274 (2022).

23 Liu, J. et al. Biological background of the genomic variations of cf-DNA in healthy individuals. Ann Oncol 30, 464–470, doi:10.1093/annonc/mdy513 (2019).

24 Hu, Y. et al. False-Positive Plasma Genotyping Due to Clonal Hematopoiesis. Clin Cancer Res 24, 4437–4443, doi:10.1158/1078-0432.Ccr-18-0143 (2018).

25 Swanton, C. et al. Prevalence of clonal hematopoiesis of indeterminate potential (CHIP) measured by an ultra-sensitive sequencing assay: Exploratory analysis of the Circulating Cancer Genome Atlas (CCGA) study. Journal of Clinical Oncology 36, 12003–12003, doi:10.1200/JCO.2018.36.15_suppl.12003 (2018).

26 Bettegowda, C. et al. Detection of circulating tumor DNA in early- and late-stage human malignancies. Sci Transl Med 6, 224ra224, doi:10.1126/scitranslmed.3007094 (2014).

27 Kabel, J. et al. Impact of Whole Genome Doubling on Detection of Circulating Tumor DNA in Colorectal Cancer. Cancers (Basel) 15, doi:10.3390/cancers15041136 (2023).

28 Yang, J. et al. Deep sequencing of circulating tumor DNA detects molecular residual disease and predicts recurrence in gastric cancer. Cell Death Dis 11, 346, doi:10.1038/s41419-020-2531-z (2020).

29 Bando, H. et al. Effects of Metastatic Sites on Circulating Tumor DNA in Patients With Metastatic Colorectal Cancer. JCO Precis Oncol 6, e2100535, doi:10.1200/po.21.00535 (2022).

30 Henriksen, T. V. et al. The effect of surgical trauma on circulating free DNA levels in cancer patients-implications for studies of circulating tumor DNA. Mol Oncol 14, 1670–1679, doi:10.1002/1878-0261.12729 (2020).

31 Reinert, T. et al. Analysis of Plasma Cell-Free DNA by Ultradeep Sequencing in Patients With Stages I to III Colorectal Cancer. JAMA Oncol 5, 1124–1131, doi:10.1001/jamaoncol.2019.0528 (2019).

32 Tie, J. et al. Circulating Tumor DNA Analyses as Markers of Recurrence Risk and Benefit of Adjuvant Therapy for Stage III Colon Cancer. JAMA Oncol 5, 1710–1717, doi:10.1001/jamaoncol.2019.3616 (2019).

## REFERENCES (Supplementary information)

1 Martin, M. Cutadapt removes adapter sequences from high-throughput sequencing reads. 2011 17, 3, doi:10.14806/ej.17.1.200 (2011).

2 Li, H. & Durbin, R. Fast and accurate short read alignment with Burrows-Wheeler transform. Bioinformatics 25, 1754–1760, doi:10.1093/bioinformatics/btp324 (2009).

3 Broad Institute, G. T. MarkDuplicates (Picard), <https://gatk.broadinstitute.org/hc/en-us/articles/360037052812-MarkDuplicates-Picard-> (2022).

4 Broad Institute, G. T. HaplotypeCaller, <https://gatk.broadinstitute.org/hc/en-us/articles/360037225632-HaplotypeCaller> (2023).

5 Broad Institute, G. T. CNNScoreVariants, <https://gatk.broadinstitute.org/hc/en-us/articles/360037226672-CNNScoreVariants> (2020).

6 Broad Institute, G. T. FilterVariantTranches, <https://gatk.broadinstitute.org/hc/en-us/articles/360040098912-FilterVariantTranches> (2020).

7 team, G. Mutect2, <https://gatk.broadinstitute.org/hc/en-us/articles/360037593851-Mutect2> (2019).

8 Kim, S. et al. Strelka2: fast and accurate calling of germline and somatic variants. Nat Methods 15, 591–594, doi:10.1038/s41592-018-0051-x (2018).

9 Christensen, M. H. et al. DREAMS: deep read-level error model for sequencing data applied to low-frequency variant calling and circulating tumor DNA detection. Genome Biol 24, 99, doi:10.1186/s13059-023-02920-1 (2023).

10 Reinert, T. et al. Analysis of circulating tumour DNA to monitor disease burden following colorectal cancer surgery. Gut 65, 625–634, doi:10.1136/gutjnl-2014-308859 (2016).

11 Pallisgaard, N., Spindler, K. L., Andersen, R. F., Brandslund, I. & Jakobsen, A. Controls to validate plasma samples for cell free DNA quantification. Clin Chim Acta 446, 141–146, doi:10.1016/j.cca.2015.04.015 (2015).

12 Forbes, S. A. et al. COSMIC: somatic cancer genetics at high-resolution. Nucleic Acids Res 45, D777–d783, doi:10.1093/nar/gkw1121 (2017).

13 Smith, T., Heger, A. & Sudbery, I. UMI-tools: modeling sequencing errors in Unique Molecular Identifiers to improve quantification accuracy. Genome Res 27, 491–499, doi:10.1101/gr.209601.116 (2017).

14 Mose, L. E., Perou, C. M. & Parker, J. S. Improved indel detection in DNA and RNA via realignment with ABRA2. Bioinformatics 35, 2966–2973, doi:10.1093/bioinformatics/btz033 (2019).

15 fgbio, f. g.-. fgbio, <http://fulcrumgenomics.github.io/fgbio/> (2021).

16 Henriksen, T. V. et al. Error Characterization and Statistical Modeling Improves Circulating Tumor DNA Detection by Droplet Digital PCR. Clin Chem 68, 657–667, doi:10.1093/clinchem/hvab274 (2022).

17 Henriksen, T. V. et al. Comparing single-target and multitarget approaches for postoperative circulating tumour DNA detection in stage II-III colorectal cancer patients. Mol Oncol 16, 3654–3665, doi:10.1002/1878-0261.13294 (2022).

